# Transport receptor occupancy in Nuclear Pore Complex mimics

**DOI:** 10.1101/2021.12.22.473839

**Authors:** Alessio Fragasso, Hendrik W. de Vries, John Andersson, Eli O. van der Sluis, Erik van der Giessen, Patrick R. Onck, Cees Dekker

**Author notes:** These authors contributed equally (Alessio Fragasso, Hendrik W. de Vries).

## Abstract

Nuclear Pore Complexes (NPCs) regulate all molecular transport between the nucleus and the cytoplasm in eukaryotic cells. Intrinsically disordered Phe-Gly nucleoporins (FG Nups) line the central conduit of NPCs to impart a selective barrier where large proteins are excluded unless bound to a transport receptor (karyopherin; Kap). Here, we assess ‘Kap-centric’ NPC models, which postulate that Kaps participate in establishing the selective barrier. We combine biomimetic nanopores, formed by tethering Nsp1 to the inner wall of a solid-state nanopore, with coarse-grained modeling to show that yeast Kap95 exhibits two populations in Nsp1-coated pores: one population that is transported across the pore in milliseconds, and a second population that is stably assembled within the FG mesh of the pore. Ionic current measurements show a conductance decrease for increasing Kap concentrations and noise data indicate an increase in rigidity of the FG-mesh. Modeling reveals an accumulation of Kap95 near the pore wall, yielding a conductance decrease. We find that Kaps only mildly affect the conformation of the Nsp1 mesh and that, even at high concentrations, Kaps only bind at most 8% of the FG-motifs in the nanopore, indicating that Kap95 occupancy is limited by steric constraints rather than by depletion of available FG-motifs. Our data provide an alternative explanation of the origin of bimodal NPC binding of Kaps, where a stable population of Kaps binds avidly to the NPC periphery, while fast transport proceeds via a central FG-rich channel through lower affinity interactions between Kaps and the cohesive domains of Nsp1.

## Introduction

Molecular traffic between nucleus and cytoplasm is exclusively controlled by the Nuclear Pore Complex (NPC), a large protein complex (52 MDa in yeast^1^) that forms a ∼40 nm-diameter pore across the nuclear envelope that encloses the nucleus^2,3^. The central channel of the NPC is filled with a meshwork of intrinsically disordered FG-Nucleoporins (FG-Nups), that feature tandem phenylalanine-glycine (FG) amino acid sequences^4,5^. Strikingly, such a FG-mesh appears to behave as a selective gate^4^ where molecules smaller than ∼40 kDa (∼5 nm in size) can pass through the pore unhindered, while the spontaneous translocation of macromolecules with size >40 kDa is strongly hampered, unless they are bound to specific nuclear transport receptors (NTRs) such as Karyopherins (Kaps) which can actively interact with, partition into, and translocate through the FG-Nup barrier. Major NTRs involved in nuclear import are Importin-β in vertebrates and its homolog Kap95 in yeast^6^.

Over the years, many models have been proposed^7,8^ to describe nuclear transport mechanistically. These can be broadly categorized into two classes: ‘FG-centric’ and ‘Kap-centric’ models. The first class of FG-centric models, which include the ‘virtual-gate’^9^, ‘selective-phase’^10,11^, and ‘forest’^5^ models, regard the barrier formed by FG-Nups as the sole important ingredient to achieve selective transport. In such a scenario, Kaps act as mere transporters, *i*.*e*., they do not take part in establishing the selective barrier. In contrast, Kap-centric models^12^, such as the ‘reduction of dimensionality’^13^, ‘reversible collapse’^14^, and ‘molecular velcro’^15^ models, predict that a part of the population of Kaps (‘slow-phase’) acts as an integral, resident component of the NPC^12,16^, while a second population of Kaps acts as transporters (‘fast-phase’). In such a two-phase model^8^, a number of Kaps (the slow-phase population) bind strongly to the FG mesh and occupy most of the available FG-repeats as a result of multivalent interactions. Notably, each Kap can bind multiple FG-repeats (up to ∼10 FG-binding sites exist in Importinβ^17^), while at the same time a single FG-Nup, which typically features several FG-repeats along its sequence, can bind multiple Kaps^18^, thus giving rise to a complex multivalent binding condensate. Saturation of available FG-repeats by the slow-phase Kaps would then result in a lowered affinity between additional free Kaps (fast-phase) and the Kap-loaded FG-mesh. Notably, such a decreased affinity would explain the occurrence of fast transit times of Kaps (∼5ms), as observed *in vivo*^19^.

Experimental evidence has been provided in support of both classes of models, and a general consensus has thus far not been achieved. A major difficulty in settling the debate stems from the limits in spatiotemporal resolution of imaging techniques^1,2,20^, combined with the complexity of the NPC in its physiological state, as it features a central mesh of ∼200 unstructured FG-Nups that are confined into a ∼40 nm pore that is constantly being crossed by many types of NTR-cargo complexes in large numbers (∼10^3^ of such protein complexes per second per pore^20^) in both directions. To probe nuclear transport through the FG-mesh, artificial mimics of the NPC have been successfully created that recapitulate the selective binding and transport behavior observed *in vivo*^21–26^. Prominent examples are biomimetic nanopores, in which ∼30-50 nm solid-state nanopores are chemically functionalized using a single type of FG-Nup (*e*.*g*. Nsp1 or Nup98) and translocations of Kaps through the reconstituted FG-mesh are monitored optically^21^ or electrically^22,27,28^. Although much knowledge has been gained in terms of a physical understanding of the FG-meshwork and its ability to impart a selective barrier, an assessment is lacking of the properties of the pore-confined FG-meshwork as a function of Kap concentration.

Here, we employ biomimetic nanopores to obtain experimental evidence on the interaction between Kap95 and Nsp1 in nanopores, with the aim to assess various aspects of Kap-centric theories of nuclear transport. Building on previous work from Ananth *et al*.^27^, which established that Nsp1-coated pores behave selectively, *i*.*e*. allowing Kap95 to pass through while blocking other inert proteins of similar size, we here investigate the behavior of Nsp1-coated pores for increasing concentrations of Kaps. Using coarse-grained modeling, we provide a microscopic view of the nanopore interior under varying Kap95 concentration, and assess the localization of Kap95 and its effect on the structural and conductive properties of the Nsp1 meshwork. To this end, we developed a coarse-grained model of Kap95 at amino-acid resolution that reproduces known binding properties between NTRs and FG-Nups.

Our data provide support for several aspects of the Kap-centric model, while also finding some discrepancies. Measurements of the ionic current through the biomimetic nanopores show fast translocations of Kaps, consistent with previous findings^27^, but on top of that, a stable shift in the baseline conductance indicates that a stable population of Kaps settles in the pore. We observe a gradual decrease in 1/f noise in the current traces as more Kaps are incorporated into the pore, consistent with a decrease in the collective fluctuations and increase in the rigidity of the Nsp1 mesh. Our simulations confirm that increasing the Kap95 concentration leads to accumulation of Kap95 near the pore wall within the nanopore, and that these Kap95 proteins have a lower mobility than Kaps located in the pore’s central channel. In contrast with predictions from Kap-centric transport theories, our modeling indicates only a slight compaction of the Nsp1-meshwork under increasing Kap95 occupation, where FG-motifs are largely unsaturated and volume limits the amount of Kap95 that can be incorporated into the pore. We attribute the existence of fast translocations on top of a stable population of Kap95 to the inherent properties of Nsp1: the extended, FG-rich anchoring domains of Nsp1 have a high avidity towards Kap95, which leads to accumulation of Kap95 near the pore wall. As the occupancy of the pore increases, additional Kap95 proteins translocate *via* a central region, formed by the cohesive domains of Nsp1, which exhibit a decreased avidity to Kap95 due to their collapsed conformation. Overall, the data show that a population of Kaps gets stably bound to the Nsp1 mesh in biomimetic nanopores and that differences in Kap95 mobility exist, supporting the idea that Kaps are an integral component of the NPC transport barrier that should be accounted for in any mechanistic model of nuclear transport.

## Results

### Conductance data show an increasing Kap95 occupancy as a function of Kap95 concentration

To perform ion current measurements through Nsp1-coated pores (Fig.1a), solid-state nanopores were fabricated in freestanding 20nm-thick SiN_x_ membranes using a transmission electron microscope (TEM, see ‘Methods’ section). Chips were mounted in a custom-built Teflon flow-cell system to allow for quick exchange of bulk solution from the two opposite compartments surrounding the chip. Conductance measurements were initially performed on freshly drilled nanopores, which, as expected, exhibited fully ohmic behavior, *i*.*e*. linear current-voltage (I-V) characteristics. Figure 1a shows the I-V plot of a bare 55 nm pore. Subsequently, pores were functionalized with Nsp1 using a 3-step self-assembled-monolayer chemistry (SAM) as described in previous work^28^ (see ‘Methods’), which yielded a ∼50% decrease in conductance (Fig. 1b). An average grafting distance of ∼6.5 nm between adjacent anchor points of the Nsp1 proteins (assuming a triangulated lattice) was estimated with surface plasmonic resonance (SPR) (Figure S3) on Nsp1-coated silica chips that were formed using the same protocol as for our nanopores.

**Figure 1.**
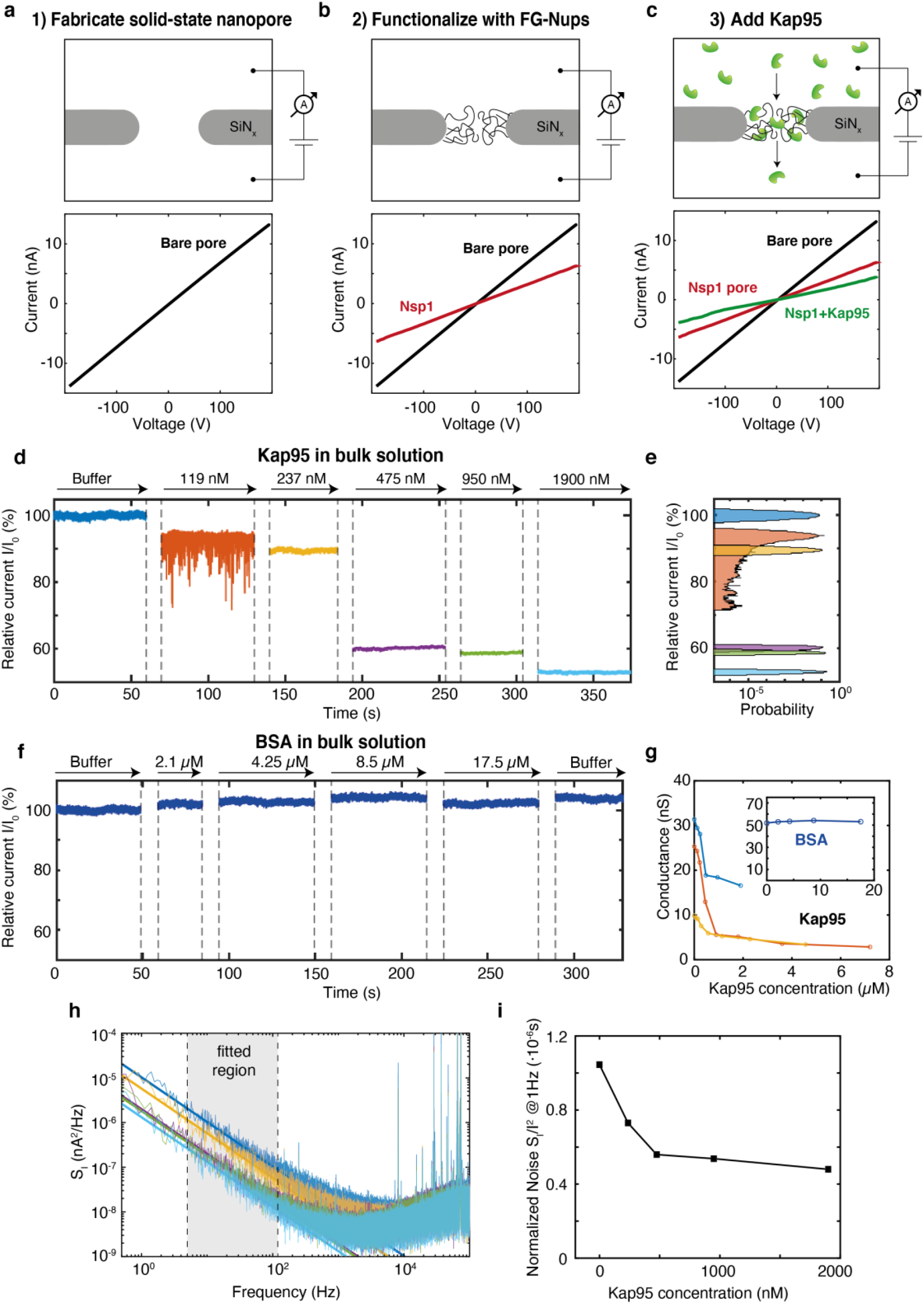
Current measurements of Nsp1-coated pores as a function of Kap95 concentration. **a**, Top: schematic showing the bare pore measurement system. Bottom: (I,V) characteristics of a bare 55 nm pore. **b-c**, Same as (**a**) but for a Nsp1-coated pore without (**b**) and with (**c**) 1.9 μM Kap95 present in the *cis*-chamber (top part). **d**, Current traces representing a Kap95 titration from 119-1900 nM, revealing a decrease of the nanopore current up to almost ∼50% of the initial value. **e**, Histogram of the current traces illustrated in (**d**). **f**, Current traces representing a BSA titration from 2-20 μM, showing no sign of current decrease. **g**, Average conductance for different nanopores *vs* Kap95 concentration. Inset: average conductance for a 55 nm pore *vs* BSA concentration (same units as in g applies). **h**, PSD spectra of the current traces shown in (**d**), same color coding as in (**d**) applies. **i**, Normalized noise power *S*_*I*_ (1 *Hz*)/*I*^2^ as a function of Kap95 concentration. The data point at 119 nM was excluded due to the presence of a pronounced Lorentzian component originating from the Kap95 translocations (see Refs.^33,73^) which yielded a poor fit of Hooge’s model to the data.

Next, yeast Kap95 was flushed on the *cis*-chamber which resulted in a further decrease of the pore conductance. This in itself directly provides a first sign that Kap95 was incorporated within the pore volume. Figure 1c (bottom) compares the three (I,V) characteristics of a bare pore (black), Nsp1-coated (red), and Nsp1-coated pore with 1.9 μM of Kap95 present in bulk solution (green), where the latter shows a further ∼50% decrease in conductance compared to the Nsp1-coated pore, indicating the presence of Kap95 molecules that interact with the pore.

To assess the pore-occupancy of Kap95 as a function of concentration, we titrated Kap95 from about 100-2000 nM (Figure 1d), and found that the pore conductance monotonously decreased in a step-wise manner as higher concentrations of Kaps were flushed into the *cis*-chamber. This clearly indicates that additional Kaps are being incorporated into the Nsp1-mesh upon increasing Kap95 concentration. Assuming that the observed conductance decrease scales linearly with the amount of Kap95 proteins present in the pore, and assuming a constant decrease of 0.80 nS due to the presence of a single Kap95 (as measured from fast Kap95 translocations through the coated pore, see next section), we estimate that ∼20 Kap95 molecules were simultaneously present in the Nsp1-pore at the highest (1.9 μM) Kap95 concentration. Interestingly, we found that transient dips in the current, caused by fast Kap95 translocation events, were present as well on top of the decreased current baseline. This effect occurred when the bulk concentration was ∼100 nM, in line with previous measurements^27^.

As a control, we repeated the same experiment by injecting increasing concentrations of BSA (Bovine Serum Albumine) in the *cis*-chamber from ∼2-20 μM (Figure 1f). Here we found that, unlike for Kap95, no significant change in the current baseline was observed. Importantly, this indicates that the interaction observed between Kap95 and Nsp1 is a result of specific protein-protein interactions, and not merely due to, *e*.*g*., electrostatic pulling of the protein into the Nsp1-mesh. Repeating the experiment on pores with a different initial conductance resulted in a similar decreasing trend of the pore conductance as a function of Kap95 concentration (Fig.1g).

Next, we analyzed the power spectral density (PSD) of the ionic current as a function of Kap95 concentration. For biomimetic nanopores, the increase in low-frequency (1-100 Hz) 1/f noise upon addition of the Nups has been typically associated to spatiotemporal fluctuations of FG-Nup mesh in the pore channel^22,27,28^. Indeed, we observed an increase in 1/f noise when comparing the bare *vs* Nsp1-coated pore (Fig.S1). Interestingly, we furthermore observed an overall decreasing trend in the 1/f noise when Kap proteins were added at increasing amounts (Figs.1h,i). To properly compare the magnitude of the 1/f noise for different current traces, we fitted the low-frequency region (5-100 Hz) of the PSD (grey area in Fig.1h) using the phenomenological Hooge model^29^, which is commonly used to describe 1/f noise in solid-state nanopores^30–33^, where the current PSD *S*_*I*_ is expressed as

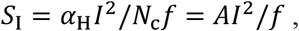

with *α*_H_ the Hooge parameter that quantifies the strength of the 1/f noise, *I* is the through-pore current, *N*_c_ is the number of charge carriers within the pore volume which depends on salt concentration and pore geometry, and *A* = *α*_H_/*N*_c_ = *S*_I_(1*Hz*)/*I*^2^ is fitted to the noise magnitude at 1 Hz normalized by the square of the current. Figure 1i shows a clear decreasing trend for *A* as a function of Kap95 concentration, indicating that a higher Kap occupancy resulted in a decrease of the 1/f noise. This reduction of the noise is suggestive of an increase in overall rigidity of the collective meshwork in the pore.

### Analysis of fast Kap95 translocations through an Nsp1 pore

We characterized the translocation events of Kap95 (fast phase) when increasing the applied voltage from 50 to 200 mV. For each event, we measured the current blockade, which to a first approximation is proportional to the volume of the translocating molecule, and dwell time, which corresponds to the time the translocating protein spends in the pore, which depends on the specific protein-pore interactions. In Figure 2a, we show examples of current traces where each current spike corresponds to a single Kap95 translocation event. Figure 2b illustrates some characteristic translocation events at different voltages in higher resolution. The asymmetric shape found in most of the translocation events (Fig.2b) may be attributed to a fast entry and association of Kap95 molecules to the Nsp1-mesh within the pore, followed by a slower dissociation and final exiting of the particle from the sensing region.

**Figure 2.**
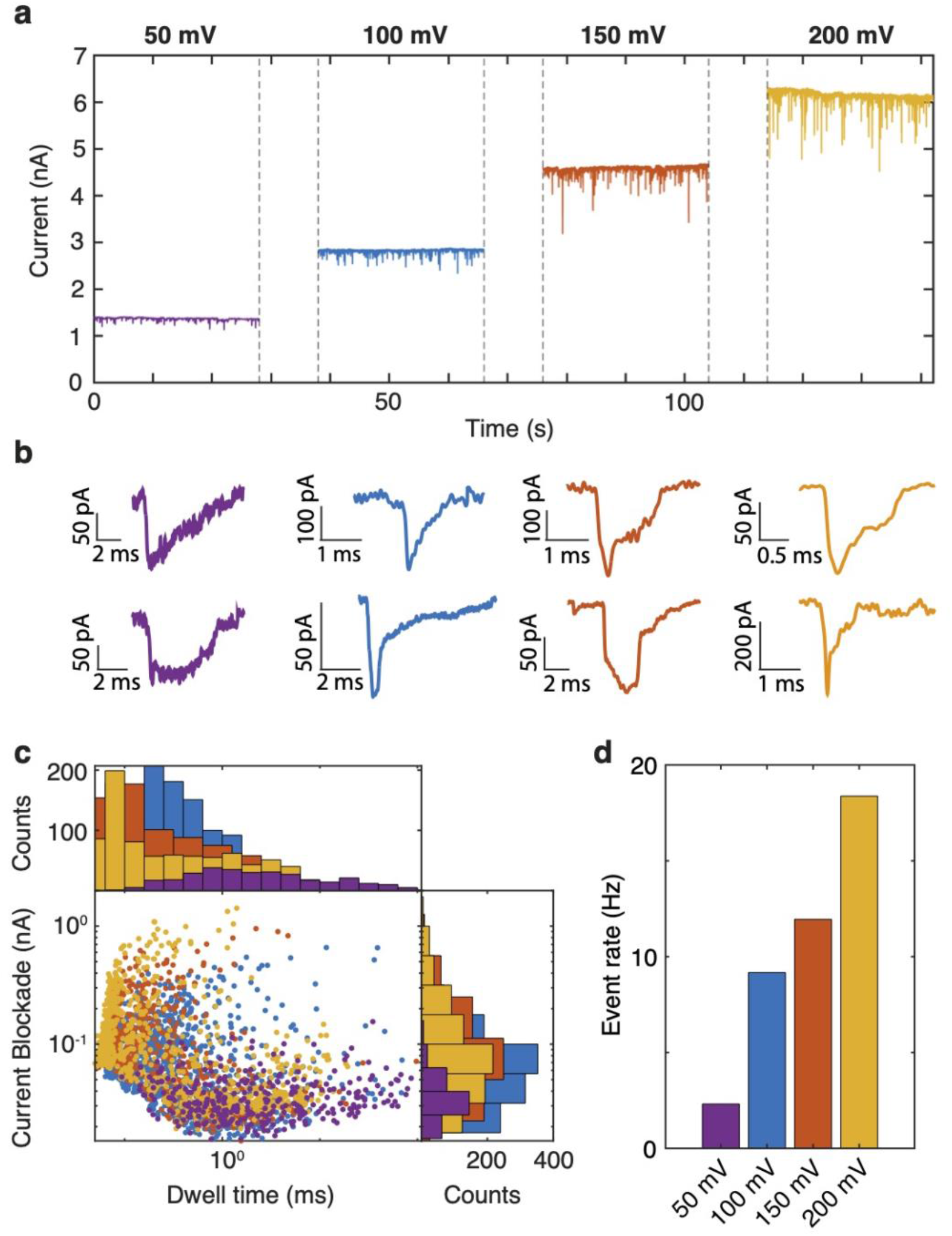
Event analysis of Kap95 translocations. **a**, Current traces showing single-molecule translocations of Kap95 through a Nsp1-coated pore under 50 mV (purple), 100 mV (blue), 150 mV (red), 200 mV (yellow). Bulk Kap95 concentration was 119 nM. Nanopore diameter was 55 nm. Additional examples are shown in Figure S2. **b**, Characteristic Kap95 translocation events at different voltages. **e**, Scatter plot of the current blockade *vs* dwell time of the translocation events for different voltages. Current blockades are taken as the maximum amplitude of the translocation event. Histograms for both dwell time and current blockade are logarithmically binned. **d**, Event rate of the translocations, defined as number of events per second, for increasing applied voltage. Color codings of b-d are the same as for **a**.

As expected, the average conductance blockades at different voltages were found to be comparable: 0.76±0.02 nS at 50 mV (N=310, errors are S.E.M.), 0.80±0.02 nS at 100 mV (N=1206), 0.77±0.03 nS at 150 mV (N=687), and 0.64±0.03 nS at 200 mV (N=820). By contrast, we observed a drastic decrease of the residence time of the protein being in the pore with increasing applied bias voltages (from 7.8±0.8 ms for 50 mV (N=310, errors are S.E.M.) to about ∼1 ms at higher voltages: 1.2±0.1 ms for 100 mV (N=1206), 0.8±0.2ms for 150 mV (N=687), and 1.1±0.1ms for 200 mV (N=820)), consistent with a stronger electrophoretic force driving the protein. Notably, the translocation events that were the least affected by the applied bias (*i*.*e*., those acquired at 50 mV) resulted in dwell times of ∼8 ms that are remarkably close to the ∼5 ms observed *in vivo*. Additional examples of current traces are shown in Fig.S2.

Lastly, the event rate of translocations, calculated as number of translocation events per second, was observed to increase as a function of applied voltage by almost an order of magnitude when increasing the bias from 50 mV to 200 mV (Fig.2d). This is indicative of an increase in the capture radius as a function of voltage^34^, defined as the radius of the hemisphere surrounding the pore wherein the electrostatic force driving the protein to the pore overtakes simple diffusion.

### Coarse-grained modeling demonstrates accumulation of Kap95 inside Nsp1-coated nanopores

To gain a microscopic view of the effects of varying Kap95 concentrations on the internal structure of Nsp1-coated pores, we performed coarse-grained molecular dynamics simulations. To this end, we developed a residue-scale model of Kap95 and combined it with an earlier-developed residue-scale computational model of Nsp1-functionalized nanopores (Figure 3a)^35,36^. In earlier works^28,35,37,38^,we employed a ‘patchy colloid’ model of Kap95, where the protein was represented by a spherical particle with the same hydrodynamic radius as the Kap95 protein, and a total of 10 binding sites modeled as hydrophobic beads. This approach was based on earlier evolutionary and computational studies^39^. Here, we present a residue-scale coarse-grained model of Kap95 that preserves the overall crystal structure of unbound Kap95 and comprises 10 FG-binding regions derived from various types of experimental^17,40,41^ data and combined computational and conservation studies^39^. Moreover, degrees of freedom that are thought to be relevant to nuclear transport, such as the charge distribution and exposed residues that participate in cation-pi interactions^42^ are included in this Kap95 model as well. The interaction strength between FG-motifs and the binding site regions is chosen such that the experimental dissociation constant of 36.1 *μ*M between unliganded yeast Kap95 (PDB: 3ND2^43^) and an Nsp1 segment ‘FSFG-K’^44^ is recapitulated (Methods, Figure S5a-c). This approach leads to multivalent and transient binding between FSFG-K and Kap95^44,45^, (Figure S5d) in line with known binding mechanisms.

**Figure 3:**
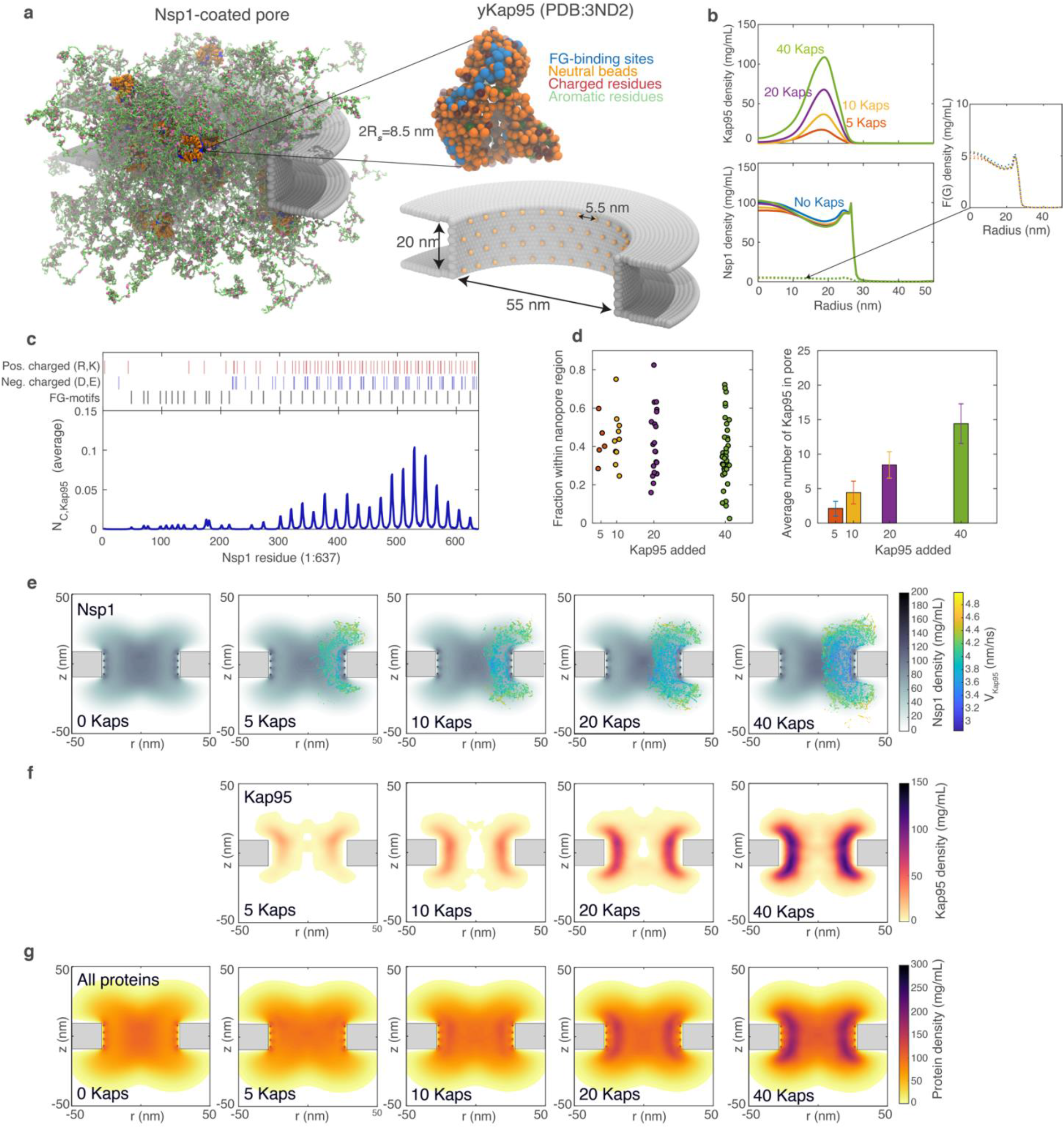
Modeling the localization of Kap95 in Nsp1-coated nanopores. **a**, One-bead-per-residue representation of Kap95 localization in an Nsp1-coated SiN nanopore. Left: Nsp1 (green)-coated nanopore, FG-motifs highlighted in pink. Right, top: single-residue model of Kap95, with essential interaction sites highlighted (blue: FG-binding site, red: charged residues, green: aromatic residues, orange: neutral beads (steric)). Right, bottom: Nsp1 anchoring geometry and scaffold structure. **b**, Kap95 and Nsp1 density profiles within the nanopore lumen (|z|<10 nm). Kap95 localizes predominantly near the pore wall, where the Nsp1 protein density is relatively low, but FG-motifs are still abundantly available. **c**, Number of contacts with a separation of <1.0 nm between Kap95 and Nsp1, averaged over simulation time, number of Kap95 molecules, and number of Nsp1 proteins. As indicated in the top panel, interactions between FG-motifs and the binding site regions on the surface of Kap95 are the main driving forces in Kap95-Nsp1 association. **d**, Probability for individual Kap95 proteins to localize within the nanopore lumen (left) and the time-averaged occupancy (number of Kap95 molecules) that reside within the nanopore lumen. Approximately 40% of the added Kap95 molecules resides inside the nanopore lumen at any given time, the remainder diffuses near the pore opening. **e**, Axi-radial, time-averaged density distributions of Nsp1 (gray scale), overlain with a velocity map of the Kap95 proteins (color scale), for varying Kap95 concentrations. The mobility of Kap95 is lower near the pore wall, an effect that becomes more pronounced at higher Kap95 concentrations. At high occupancies, Kap95 molecules that reside close to the pore wall are on average 50% less mobile than Kap95 molecules diffusing near the central channel and pore opening. **f**, Axi-radial and time-averaged density distribution of Kap95 showing that Kap95 preferably localizes in a band near the pore wall. **g**, Axi-radial and time-averaged density distribution of the cumulative protein mass (Kap95 and Nsp1) inside the nanopore. In absence of Kap95, the lowest protein concentration is found near the pore wall, whereas for increasing Kap95 occupancy, the lowest protein concentration is found in the central channel of the pore.

We then performed coarse-grained molecular dynamics simulations of Kap95 localization in Nsp1-coated nanopores using a one-bead-per-amino acid model for disordered FG-Nups^46,47^ and our newly-parametrized model for Kap95 (Methods). Nsp1-proteins were end-grafted to the interior wall of a nanopore occlusion with a diameter of 55 nm and a thickness of 20 nm (Figure 3a), closely approaching the values used in our experiments (Methods). We varied the number of Kap95 proteins between 0 and 40 (Figure 3a). The number of Kap95 molecules in the pore region would, if spread out evenly over the simulation volume, correspond to a wide range of concentrations from 0 to ∼10^2^ μM.

The time-averaged density distributions that we obtained for the Kap95 proteins (Figure 3b, top and 3g) demonstrate that, for all Kap95 concentrations, Kap95 predominantly localizes peripherally (*i*.*e*., near the pore wall). This finding can be explained by considering the distribution of Nsp1 protein mass and FG-residues (Figure 3b, bottom). Large molecules such as Kap95 were observed to preferably localize in the sparse regions of the Nsp1-coated pore where FG-motifs are still abundantly available but where the steric hindrance is lowest. By means of a contact analysis, we found that Kap95 and Nsp1 predominantly associate by virtue of binding between FG-motifs in the highly-charged Nsp1 domain (Figure 3c) and the binding pockets on the surface of Kap95 (Figure S7), a finding consistent with the association mechanism deduced in NMR measurements^44^. The peripheral localization of Kap95 is consistent with other computational studies on a planar geometry, which highlighted that model Kap95 particles, by virtue of their large size, preferably localize in sparse regions in Nsp1 brushes^48^.

We calculated the probability for an individual Kap95 protein to enter the Nsp1 nanopore lumen as well as the corresponding average number of Kap95 molecules present within the pore lumen. Our results in Figure 3d (left panel) indicate that there is a significant spread in the probability for a single Kap95 molecule to reside in the pore meshwork. The average occupancy of the pore lumen was found to be approximately 40% (Figure 3d), a finding that was found to hold for all local concentrations of Kap95. For 20 Kaps in the pore, the conductance was reduced by 45 % (Fig. 4d), a number that nicely corresponds to the experimental observations at 1.9 μM (Fig.1d).

**Figure 4:**
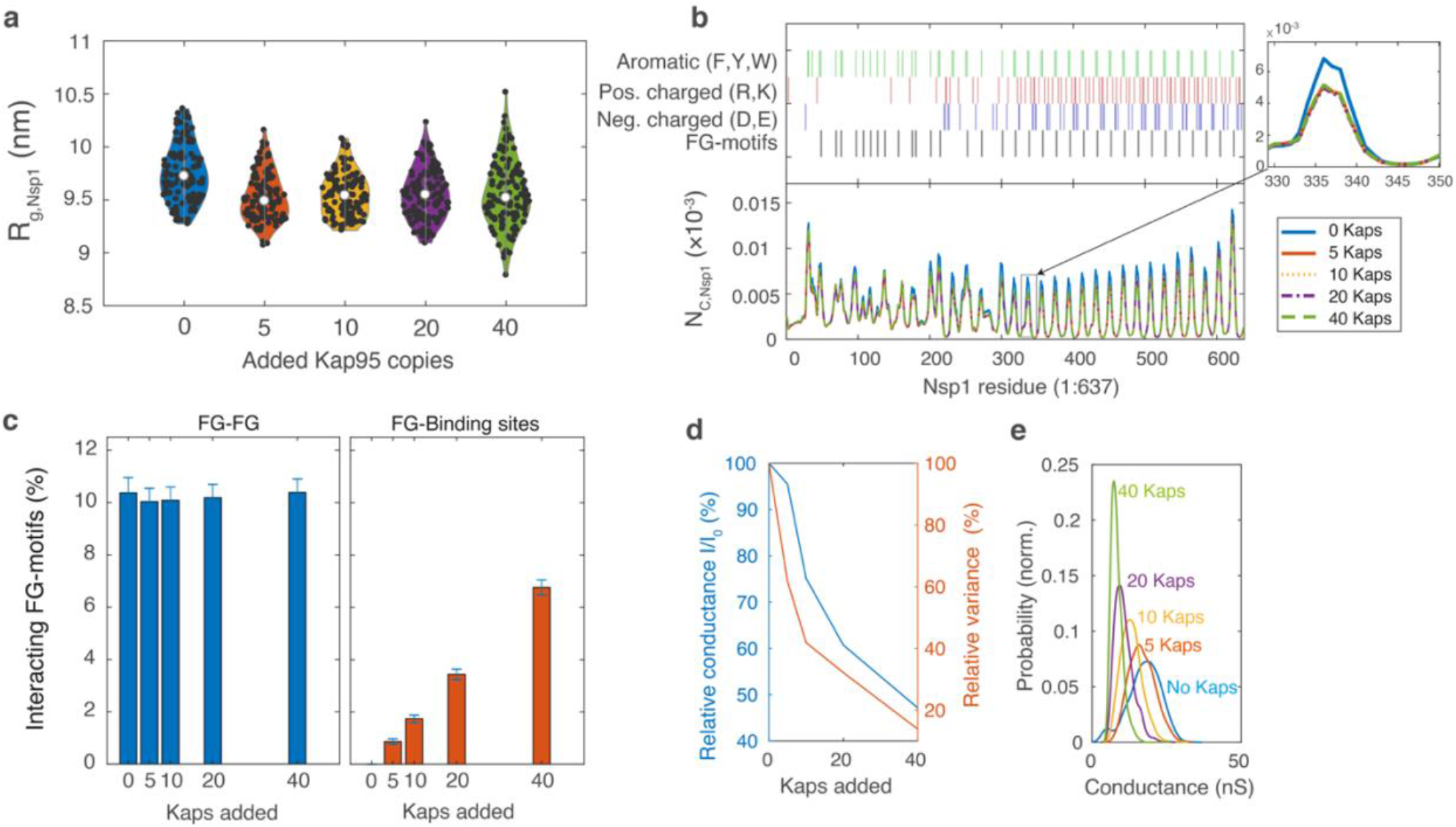
Effect of Kap95 concentration on the properties of the Nsp1-coated nanopore. **a**, Violin plot of the Nsp1 radii of gyration (N=120). The Nsp1 molecules appear to exhibit a slight compaction upon addition of Kap95. **b**, Contacts between Nsp1-proteins (averaged over time and the number of proteins) as a function of residue index (bottom panel), where specific residue types are highlighted (top panel). Intra-Nsp1-contacts mainly take place via FG-motifs in the extended region (173-637), or via hydrophobic/polar contacts in the cohesive domain (1-172). Inset: Zoom of 1 peak. Addition of 5 Kap95 molecules induces a slight (<10 %) decrease in intra-Nsp1 contacts in the extended domain, but we observe no appreciable change in the intra-Nsp1 contacts for further increases in Kap95 concentration, leading to overlapping curves for non-zero Kap95 concentrations. **c**, Left panel: average (± S.D.) number of inter-FG-motif contacts (‘FG-FG’) within the Nsp1 meshwork, which appears to be nondependent on the Kap95 concentration. Right panel: the number of FG-motifs involved in Kap95 binding (‘FG-BS’) scales linearly with the number of Kap95 molecules in the nanopore. With only maximum ∼7% of the -motifs involved in Kap95 binding, the total number of unbound FG-motifs remains high in all cases, indicating that only a small fraction of the FG-motifs is occupied. **d**, Left axis: Mean conductance as a function of added Kap95, normalized by the conductance of the Nsp1-coated nanopore with no Kap95 present. Similar to our experiments, we find that the addition of a small number of Kap95 molecules strongly reduces the average nanopore conductance. Right axis: the variance in the conductance value decreases with increasing Kap95 concentration. Values were calculated from 10 ns simulation windows. **e**, Probability distribution of the nanopore conductance for different numbers of added Kap95 molecules. Similar to our experimental results, d-e highlight that an increasing Kap95 occupancy leads to a decrease in conductance and a reduction in the current fluctuations.

Interestingly, the center-of-mass displacements of Kap95, calculated on a longer timescale and superimposed onto the axiradial density distribution of Nsp1 (Figure 3f), demonstrated that the mobility of Kap95 inside the pore region depends on the concentration of Kap95. Whereas for low Kap95 concentrations (*e*.*g*., for 5 or 10 Kap95 molecules), the spatial variation in Kap95 mobility was found to be relatively small, higher concentrations of Kap95 led to more pronounced spatial differences in mobility. Consistent with predictions by Kap-centric transport theories, we found that, at high concentrations, Kap95 molecules residing near the pore wall had a 50% lower mobility than Kap95 residing near the center of the channel.

### Kap95 affects the structural properties of Nsp1-coated nanopores in a concentration-dependent manner

Kap-centric transport theories^8,16,49^ suggest that the presence of NTRs can affect the conformations of FG-Nups within assemblies and that the multivalent binding interactions between NTRs and FG-Nups leads to an effective saturation of the FG-meshwork. Both of these statements are mainly deduced from findings on planar assemblies of FG-Nups^15,50^, although evidence also exists that the presence of Kap95 homologs may lead to a swelling and stiffening of the interior of the NPC^51^. To assess the effect of Kap95 on the internal structure of Nsp1-coated pores, and to gain a microscopic view of the conductance processes in our experiments (Figs. 1d-i, 2), we calculated various quantities that describe the Nsp1 meshwork as a function of local Kap95 concentration. More specifically, we assessed the changes in Nsp1 single-molecule properties, FG-motif binding characteristics, and more macroscopic properties of the nanopore system such as the ion conductance.

The presence of Kap95 induces a small but clearly observable compaction of Nsp1, evidenced by a ∼5 % decrease in the radius of gyration of Nsp1 (Figure 4a) and its average persistence length (Figure S8). Earlier QCM-D and ellipsometry experiments^50,52^ suggested different responses of Nsp1-coated surfaces to Kap95 titration, *e*.*g*. a compaction followed by swelling, or only slight swelling. The small effect of Kap95 on Nsp1 morphology in our simulations, even at high concentrations, differs from certain predictions of Kap-centric transport models such as large conformational changes or the opening of a central transport channel.

To assess whether Kap95 can compete with Nsp1 for FG-motifs, we calculated the fraction of FG-motifs that interact with other FG-motifs or with Kap95 binding sites (Figure 4c). Expressed as the fraction of total FG-motifs present inside the nanopore system, we find that, regardless of Kap95 concentration, approximately 10.5% of the FG-motifs are in contact with another FG-motif. The fraction of FG-motifs that interact with binding site regions on the Kap95 molecules scales linearly with the number of Kap95 molecules added, and is at most ∼7 % for the highest local Kap95 concentration studied here (40 Kaps; ∼10^2^ μM). Based on the average number of FG-binding site contacts for each Kap95 concentration (Figure 4c, right panel), we find that a single Kap95 protein interacts with up to 7 FG-motifs simultaneously. The large fraction of unbound FG-motifs at any point in time indicates that the Nsp1-meshwork is not saturated in terms of available binding sites: considering an average valency of 7 FG-motifs bound per Kap95 molecule, a local concentration of ∼500 Kap95 molecules per pore would be required for complete saturation, a number that we consider unfeasible given the finite unoccupied volume of the pore lumen. We therefore conclude that, interestingly, there is no strong competition between Kap95 and Nsp1 molecules for FG-motifs.

To understand the effect of Kap95 concentration on baseline ion conductance, the conductance blockade caused by Kap95, and the PSD observed in our experiments (Figure 1d-e,g), we calculated the conductance of Nsp1-coated pores under varying Kap95 occupancies. In line with earlier work^28,35,36^, we employed a modified Hall equation^53^ (see Methods), where we assumed a linear dependency of the local conductivity on average protein density, up to a cutoff density. We found that increasing the local Kap95 concentration mainly affected the density at larger radial coordinates (Figure 3b, top panel), which were originally the sparsest (Figure 3b, bottom panel), and therefore most strongly conducting regions. The local protein concentration (i.e. the sum of Kap and Nsp1 concentrations) already exceeded the cutoff value in our conductance model at relatively low Kap95 occupancy, leading to a strong conductance decrease at small Kap concentrations (Fig. 4d). The decrease in baseline conductance became less pronounced with increasing Kap95 concentration, consistent with the trends in Figure 1d. Like in the experiments, a decrease in the baseline ionic conductance of ∼ factor 2 was observed. Moreover, the noise in the conductance values decreased notably, as is clear from the reduced variance in the conductance (Figure 4d) and the reduced width of the conductance probability density distribution (Figure 4e). This finding also qualitatively agrees well with the conductance results in Figure 1i, where we found that the 1/f noise reduced with increasing Kap95 occupancy. The absence of measurable Kap95 translocation events at high concentrations (Figs. 1d,e) may be explained through our density-based conductance analysis: the effect of a Kap95 translocation on the local conductance diminishes as the regions near the pore wall, which were originally sparse and therefore highly conductive, are now increasingly populated. Subsequent translocations of Kap95 molecules occur through the dense regions of the central cohesive conduit that is less conducting, and these Kaps will therefore exhibit a smaller conductance blockade.

In summary, our simulations indicate the presence of Kap95 in the peripheral region within the nanopore lumen which only slightly affected the morphology of the Nsp1-meshwork. Moreover, the total amount of available binding sites (FG-motifs) for Kap95 in the Nsp1-meshwork is very large and does not diminish notably, even for very high local Kap95 concentrations. This indicates that the number of Kap95 proteins that can be harbored by the Nsp1-nanopore is limited by the finite pore volume rather than a reduction in available binding sites. We find that the stable drop in the conductance baseline upon flushing increasing Kap95 concentrations is a consequence of a stable population of Kap95 occupying the peripheral regions of Nsp1-coated pores.

### Consequences of these findings for Kap-centric models of nuclear transport

It is of interest to discuss these new experimental and modelling data in the context of the various Kap-centric models for nuclear transport. Most importantly, our experimental nanopore data show that while Kaps are rapidly translocating through the pore in a timescale of milliseconds, a second population of quasi-permanent Kaps is also present. Such slow population appears in the ion current data as a stable, concentration-dependent decrease of the current baseline. Our computational modeling shows that Kap95 localizes near the pore wall, which sharply decreases the conductance with increasing Kap95-concentration in agreement with the experimental data. Moreover, binding of Kap95 appears to have a ‘stiffening’ effect on the Nsp1-nanopore interior as a whole, as observed by the overall decrease of the 1/f noise in the nanopore. Our density-based conductance model as confirms that increasing Kap95 occupation reduces the variability in the conductance, consistent with our observations of reduced 1/f noise.

Kapinos *et al*.^12^ suggested a possible arrangement of the two Kap95 populations (fast *vs* slow) inside the nanopore where slow Kaps that bind the FG-Nup mesh with high avidity would induce a collapse of the mesh towards the pore rim, resulting in the opening of a central channel through which fast Kaps can rapidly diffuse through at ∼ms time scales. Instead, and contrary to this prediction of the Kap-centric model, our simulations highlight that the Nsp1-meshwork does not collapse in the presence of Kap95 to accommodate a central opening or exposed surface. We do see, however, a stable population of Kap95 molecules does exist near the pore wall, which is less mobile than the Kap95 molecules diffusing through the pore center. We can attribute this to initial Kap95 having a greater affinity for the extended anchoring domains of Nsp1 (Figure 3c), where the steric hindrance is lower (Figure 3b, bottom panel) and where the exposed FG-motifs are more readily accessible compared to the centrally located cohesive head groups (Figure 5b). This naturally leads to a spatial segregation of the Kaps in the nanopore and the coexistence of slow and fast populations (Figures 1d, 3e-g, 5). Namely, as the anchoring regions of Nsp1 near the pore wall are increasingly occupied by Kap95 (Figure 3g), translocating Kaps may instead prefer to traverse via the central region, where the total protein density and thus steric hindrance is lowest (Figure 3f,g) and where cohesive Nsp1 head regions are present with a lower affinity for Kap95 (since less FG-motifs are exposed, Figure 5b) and thus a higher mobility (Figure 3e). As two populations of Kap95 arise without the need of an exposed surface, our results form an alternative to the ‘reduction-of-dimensionality’^13^ and ‘molecular velcro’^15^ models, where translocating Kaps would cross the pore following a 2D-random walk over a collapsed, Kap-saturated layer of FG-Nups.

**Figure 5:**
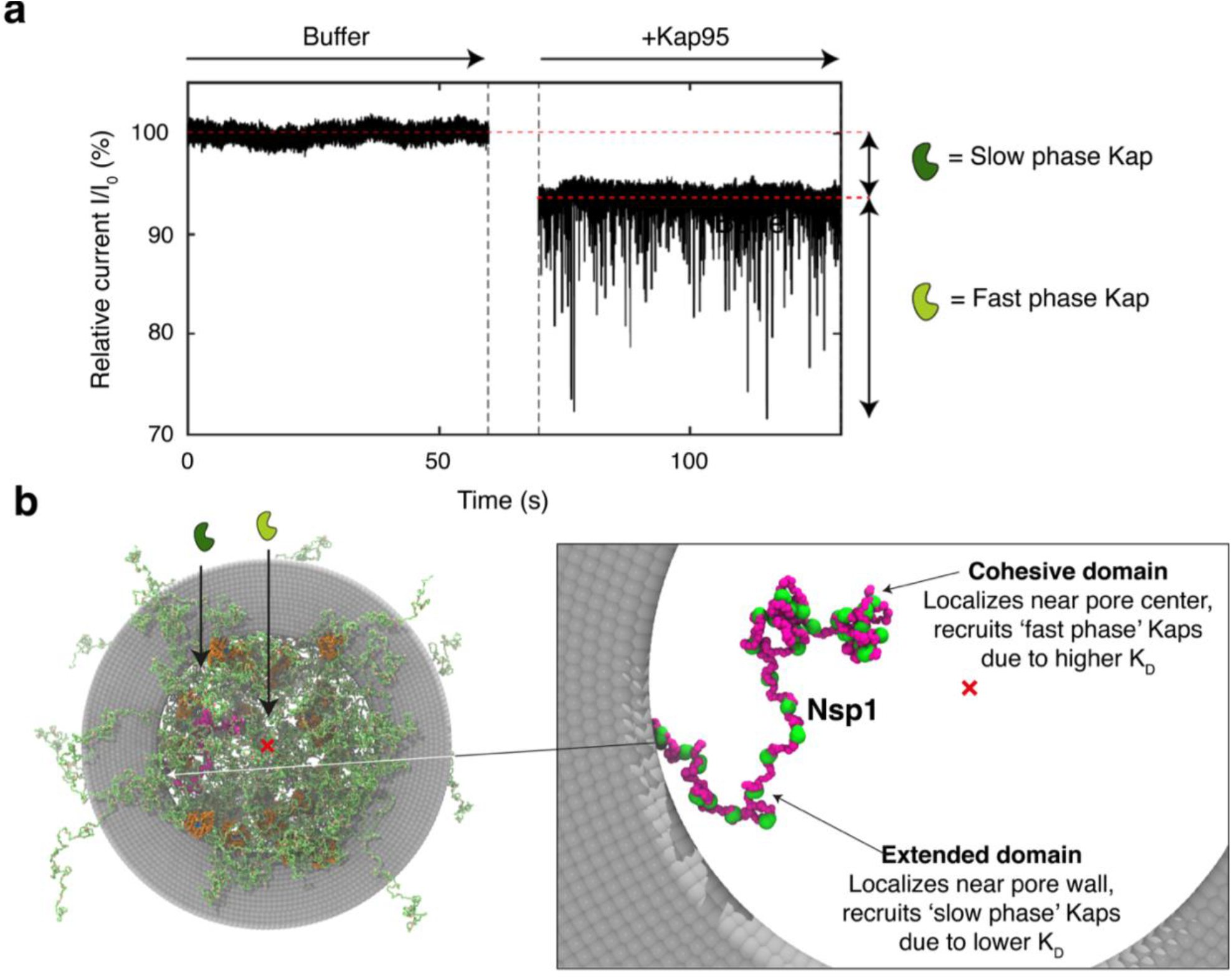
Kap-centric transport mechanism in Nsp1-coated pores. **a**, Nanopore current traces, where a decrease in baseline is associated with a ‘slow phase’ of Kaps, which are stably incorporated in the Nsp1 mesh, and transient spikes probe the ‘fast phase’ of Kaps that translocate on ∼ms timescales. **b**, Slow phase Kaps localize in the low-density region near the pore wall by interacting with the extended and FG-rich anchoring domain of Nsp1. A single copy of Nsp1 is highlighted in purple for visual illustration. Red cross indicates the nanopore center. Right image provides a zoom of the highlighted single Nsp1-protein with its two-domain structure. The FG-motifs (green) are exposed leading to high avidity with Kap95. As the anchoring regions are increasingly occupied, a fast phase of Kaps is forced to interact with the cohesive headgroups of Nsp1, that assemble in the central region of the Nsp1 pore and interact less strongly with Kaps. Due to the two-domain structure of Nsp1 with different avidities towards Kaps, a spatial segregation arises naturally.

Moreover, we observe that the Kaps that enter the Nsp1 meshwork only occupy a small fraction of the available FG motifs (less than 8% at the highest Kap concentration, see Figure 4c). Since the fraction of FG-motifs that partakes in intra-FG contacts also remains small (∼10.5% for all Kap95 concentrations, see Figure 4c), the number of free FG-motifs far exceeds the sum of Kap95-bound and FG-bound FG-motifs, meaning that the Nsp1-meshwork is not saturated in terms of available FG-motifs. Instead, our results suggest that the FG-motifs become sterically unavailable due to the high occupancy of Kaps that fill up the pore.

It is important to place our results on Kap occupancy in Nsp1 nanopores into the context of the complexity of the entire NPC. As was demonstrated in the original studies on Kap-centric transport theories^25,49,54^, both the anchoring geometry (*e*.*g*., grafting density) and physicochemical properties (*e*.*g*., number and type of FG-motifs and spacer cohesivity) of FG-Nups control the response of an FG-Nup assembly to increasing concentrations of Kap95. In contrast to essential yeast FG-Nups that have a bimodal (highly-charged versus cohesive) structure such as Nup100, Nup116, and Nup145N, the highly charged anchoring domain of Nsp1 contains a large number of FG-motifs. The lack of strong conformational change in Nsp1 upon increasing presence of Kap95 may be largely explained by the association of Kap95 with these extended, FG-rich regions. To further assess the validity of Kap-centric transport models, future studies could additionally assess the transport properties and morphology of biomimetic nanopores that comprise FG-Nups that carry no FG-motifs in the extended domain, or FG-Nups that lack a two-domain structure entirely: changes in conformation of such FG-Nups with increasing Kap95 concentrations may be more pronounced than for the Nsp1 protein studied here.

## Conclusion

In this work, we investigated the interaction between the yeast transporter Kap95 and FG-nucleoporin Nsp1 using biomimetic nanopores and coarse-grained modeling. We identified two distinct Kap95 populations: slow Kaps that bind stably to the Nsp1 mesh causing a higher protein density and according decrease in the current baseline, and fast Kaps that rapidly translocate the pore on a ∼ms timescale. Our simulations identified that the stable population of Kap95 resides near the pore wall by associating with the extended, FG-motif-containing anchoring regions of Nsp1. The population of slow Kaps was found to increase in a concentration-dependent manner as revealed by the strong decrease of the pore conductance, consistent with the modeled structure of the Nsp1-Kap95-meshwork. The center of the pore harbors a larger number of fast Kaps than the region near the pore wall, where the higher mobility was associated with a lower total protein density and a lower availability of exposed FG-motifs compared to the densely populated region near the pore wall.

Taken together, our data agree with some, but not all predictions of Kap-centric nuclear transport models. In brief, we find that Nsp1-coated pores show, as predicted, fast ∼ms transport and a stable population of Kap95 (which localizes near the pore wall), but we did not observe a significant Kap95-induced collapse in the Nsp1 meshwork. Notably our experiments were carried out on 1 type of FG-Nups, whereas the full NPC contains ∼12 different types FG-Nups, each present in a well-defined stoichiometry and positioning along the NPC lumen.

The approach presented in this work can be expanded in many ways. For example, it paves the way to studies that mimic more physiological scenarios by, *e*.*g*., involving multiple types of Nups, introducing the Kap95 binding partner, Kap60, and a RanGTP-assisted dissociation when RanGTP is added on the *trans*-side of the pore. The approach towards modeling of Kap95-FG-Nup interactions can be applied to other proteins from the Karyopherin family, opening up the possibility to study the effect of conformational changes or cooperativity on Kap-FG-Nup binding from a computational perspective. Other platforms that avoid the use of an applied voltage can also be envisioned, *e*.*g*. by using zero-mode-waveguides (ZMW^55^), where translocating molecules are freely diffusing (*i*.*e*. not driven by an electrical field) and optically detected. Finally, probing the spatial localization of Kap95 within Nsp1-coated pores, here done *in silico*, can in principle also be studied experimentally by labelling Kap95 with gold nanoparticles and performing CryoEM imaging of the pores. More is to come.

## Acknowledgements

We would like to thank the Görlich Lab for sharing purified Nsp1, Meng-yue Wu for technical assistance on the TEM, and Marc op den Kamp, Sjoerd Meesters and Koen Wortelboer for their assistance in developing precursors to the Kap95 model. This research was funded by NWO-I programme ‘Projectruimte’, grant no. 16PR3242-1. We acknowledge discussions with Nils Klughammer, Paola de Magistris, Anders Barth, Adithya Ananth, Sonja Schmid, Hamid Jafarinia and Mark Driver. H.W.d.V. acknowledges support from the CIT of the University of Groningen and the Berendsen Centre for Multiscale Modeling for providing access to the Peregrine and Nieuwpoort high performance computing clusters. C.D. acknowledges support from the ERC Advanced Grant no. 883684 and the NanoFront and BaSyC programmes.

## Author contributions

A.F. and C.D. devised the experiments. H.W.d.V, E.v.d.G., and P.R.O. devised the simulations.

E.O.v.d.S. cloned and purified the proteins. A.F. carried out the nanopore experiments and analysis.

J.A. carried out the SPR experiments and analysis. H.W.d.V. carried out the simulations and analysis. A.F., H.W.d.V., P.R.O. and C.D. wrote the manuscript.

## Data availability

Data that support the findings of this study are available from the corresponding authors upon reasonable request.

## Methods

### Preparation of solid-state nanopores

Pores with sizes ∼40-55 nm were fabricated onto freestanding 20 nm-thick SiN_x_ membranes supported on a glass substrate for low-noise recordings (purchased from Goeppert). Drilling of the pores was performed by means of a transmission electron microscope (see Ref.^56^ for details). Pore functionalization was performed as reported previously^28^. Briefly, freshly drilled nanopores were rinsed in milliQ water, ethanol, acetone, isopropanol, and treated with oxygen plasma for 2 min to further clean the chip and enrich the nanopore surface with hydroxyl (-OH) groups. Next, the chip was incubated with 2% APTES (Sigma Aldrich) in anhydrous toluene (Sigma Aldrich) for 45 min, at room temperature, shaking at 400rpm, in a glove-box filled with pure nitrogen, which prevented APTES molecules from polymerizing. The chip was subsequently rinsed in anhydrous toluene, milliQ water, and ethanol, blow-dried with nitrogen, and heated at 110°C for ∼30-60min. Following the curing, the chip was incubated with Sulfo-SMCC (sulphosuccinimidyl-4-(N-maleimidomethyl)-cyclohexane-1-carboxylate) (2 mg no-weight capsules (Pierce)) for >3hrs, a bifunctional crosslinker that binds amine groups of the APTES through a NHS-ester group, while providing a free maleimide group on the other end. Chips were then rinsed in PBS for 15 min and incubated with Nsp1 for ∼1 hr, which reacted with the maleimides through the cysteine present on its C-terminus, forming stable covalent bonds. The chips were finally rinsed with PBS to remove unspecifically bound proteins.

### Electrical ion-current measurements

The buffer employed in nanopore experiments was 150 mM KCl, 10mM Tris, 1mM EDTA, pH 7.4. Current data were recorded in real-time using a commercial amplifier (Axopatch200B, Molecular devices), which applies a 100kHz low-pass filtering, and digitized at 250kHz (Digidata 1322A DAQ). Raw traces were further filtered digitally at 5kHz, and processed using a custom-written Matlab script^57^.

### Purification of Kap95

We refer to Ref.28 for details on the purification of Kap95.

### SPR measurements and analysis

SPR measurements were performed in air using a Bionavis MP-SPR Navi^Tm^ 220A instrument equipped with two 670 nm laser diodes directed on two different spots on the sample surface. Silicon dioxide coated SPR sensor slides were prepared from regenerated gold sensors or glass substrates (Bionavis), according to the following procedure. Substrate cleaning and removal of previous metal layers was performed using RCA2 treatment (HCl:H2O2:MQ-water at 1:1:5 volume ratios at 80 °C for 30 min) followed by O_2_-plasma treatment at 50 W, 250 mTorr, 80 sccm for 60 seconds. Metal deposition of 2 nm Cr and 50 nm Au was then performed using electron beam physical vapor deposition (Lesker PVD 225) and final deposition of SiO_2_ was performed with atomic layer deposition (Oxford FlexAL) at 300 °C using a BTBAS precursor and O_2_ as process gas. The optical background of all SPR sensors was measured individually prior to being coated in the procedure described under “‘Preparation of solid-state nanopores”. Furthermore, material loss corresponding to 1.1 nm of the SiO_2_ layer was measured on a reference sensor following the cleaning with O_2_-plasma as described in the previous section. The model background of all samples was corrected with the same amount accordingly. The sensors were briefly rinsed with ethanol (Nsp1 samples were instead rinsed in MilliQ-water) and blow dried in a gas stream of N_2_ immediately before measurements. Adlayer thickness and grafting distance was determined from least-square fitting measurements with Fresnel models as described previously^28,58,59^, with optical parameters given in Table 1 below. When calculating grafting density and grafting distance of Nsp1 (*M*_Nsp1_ = 65.7 kDa) a density of 1.35 g/cm^3^ was assumed^60^.

**Table 1.** Optical parameters for each layer included in the Fresnel models. The values enclosed in brackets were incorporated in the model background for subsequent layer(s).

|  | **Prism** | **Cr** | **Au** | **SiO<sub>2</sub>** | **APTES** | **SMCC** | **Nsp1** |
| --- | --- | --- | --- | --- | --- | --- | --- |
| **d (nm)** | - | 2 | 50 | ~ 9.5 | Fitted (2.3) | Fitted (2.6) | Fitted |
| **n** | 1.5202 | 3.3105 | 0.2238 | 1.4628<sup>61</sup> | 1.4200 | 1.4200* | 1.5300<sup>62</sup> |
| **k** | 0 | 3.4556 | 3.9259 | 0 | 0 | 0 | 0 |

\*Value unknown. The given value is used for comparison with APTES.

### Coarse-grained model for unfolded proteins

The coarse-grained molecular dynamics simulations in this work are all performed using a modified version of the implicit-solvent one-bead-per-amino acid (1-BPA) model for unfolded proteins developed and applied earlier^27,36–38,46,47,63^. The 1-BPA model accounts for the physicochemical properties (charge, hydrophobicity) of all 20 amino acids, and incorporates backbone potentials that distinguish between three groups of amino acids (*i*.*e*., Glycine, Proline or other residues) depending on their backbone stiffness. The interaction potentials between cationic residues (R,K) and aromatic residues (F,Y,W) have been recalibrated to accommodate for the effect of cation-pi interactions^64^. We refer the reader to Refs. ^46,47^ and section 4 of the Supporting Information for a more in-depth discussion of the forcefield, and recapitulate the non-bonded potentials and cation-pi interaction parameters in Tables S1-4 and Equations S1-4 for completeness. All simulations were carried out at a temperature of 300K and timestep of 15 or 20 fs (Table S6), where the use of Langevin dynamics accounts for solvent viscosity and implicitly facilitates temperature coupling. An inverse friction coefficient *τ*_T_ of 50 ps was used. All simulations were performed using the GROMACS software suite^65,66^ (versions 2018.4 and/or 2016.3) on a parallelized computing cluster.

### Parametrization of a coarse-grained model for yeast Kap95

We based the residue-scale coarse-grained model for yeast Kap95 on the unbound crystal structure (PDB ID: 3ND2^43^), where beads representing single amino acids were centered on the corresponding alpha carbon positions. The secondary and tertiary structure of Kap95 were preserved using an elastic network: for all amino acid beads separated less than 1.2 nm, a harmonic potential with a binding constant of 8000 kJ/nm^2^ was applied, with the original separation between amino acids beads in the crystal structure forming the reference distance for the harmonic potentials. Within the 1-BPA-CP (‘cation-pi’) forcefield^64^, interactions between Kap95 and unfolded proteins fall under one of four categories (see sections 4 and 5 of the SI, Tables S3-4): coulombic interactions, volume exclusion, cation-pi interactions and specific interactions between binding site regions and FG-residues that use the 1-BPA hydrophobic potential (Equation S1). We assigned Kap95 residues to binding site regions based on the following procedure: First, we considered all binding site residues identified in a computational study^39^ that investigated the binding of FG-Nup segments and isolated FG/GLFG-repeats to evolutionarily conserved regions on the surface of a mouse homolog of Kap95 (PDB: 1UKL^67^). Based on a sequence alignment^68,69^ we then identified corresponding residues in yeast Kap95 (Table S5). Visual inspection and structural alignment using VMD^70^ confirmed that binding sites were spatially well-conserved (Figure S4). In order to derive an accurate value for the interaction parameter between binding site regions and FG-motifs, *ϵ*_BS,FG_, we calculated the dissociation constant *K*_D_ between Kap95 and a highly charged Nsp1-segment termed FSFG-K (Figure S5a-c), and assessed the number and duration of contacts between FG-motifs and binding sites (Figure S5d).

The dissociation constant *K*_D_ between Kap95 and FSFG-K was calculated using the following relation^71^:

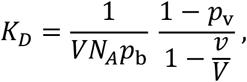

where *p*_b_ is the fraction of bound configurations with the minimum distance *d*_*ij*_ < 0.8 nm^71^, *p*_v_ the fraction of configurations where the center of mass of FSFG-K resides in the sub-volume *V*_sub_ centered around Kap95, *N*_A_ Avogadro’s number and *V* the box volume (fixed at 20^3^ nm^3^ throughout). The sub-volume *V*_sub_ is defined as a spherical volume centered around the center-of-mass of Kap95 with radius 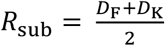, where *D*_F_ and *D*_K_ are the average largest diameters (largest internal distance between any residue) of FSFG-K and Kap95, respectively. The number of contacts in the calculation of *p*_b_ and the contacts between individual binding sites and FG-motifs were determined using the MDAnalysis Python package^72^, version 1.9.

New paragraph. Since the number of residues per binding site, as well as their exposedness to FG-motifs at the Kap95 surface varied between binding sites, additional residues were added to binding sites 1,5,6 and 10 (Table S5). This reduced the differences in contacts between binding site residues. Given that the experimental *K*_D_ -value is considered a lower limit^44^ and that we observed transient binding behavior for a range of *ϵ*_BS,FG_-values, we performed additional simulations as a secondary verification (see section 5 in the Supporting Information for additional details): We inserted 10 copies of Kap95 in a 30 nm diameter biomimetic nanopore coated with an artificial FG-Nup ‘NupX’. In earlier work, we demonstrated that Kap95 is able to translocate through such pores^28^. We identified an optimum value of *ϵ*_BS,FG_ = 13.75 kJ/mol (Figure S6) based on the criterion of Kap95 remaining in the NupX meshwork while maintaining transient contacts with the FG/GFLG-motifs in NupX. This value is used throughout the remainder of this work.

Nsp1-coated nanopores were modeled as follows: a cylindrical occlusion with a diameter of 55nm is generated, consisting of sterically inert particles with a diameter of 3 nm. Nsp1 is then anchored to the interior of the nanopore by its C-terminal Cys-residue, in four rows in a triangulated fashion, using a grafting distance of 5.5 nm (see Fig. 3a). This value is based upon estimates from earlier work^35^ and slightly smaller than estimates (which are likely to slightly overestimate the grafting distance due to dehydration and rehydration steps) from SPR measurements in the current work. A similar procedure is used for the NupX-coated nanopores, where a diameter of 30 nm is used and NupX is also anchored by its C-terminal Cys-residue. Kap95 molecules were added in either one (5, 10 copies, *cis*-side), two (20 copies, *cis*-side) or four (40 copies, *cis*- and *trans*-side) layers and pulled into the Nsp1 meshwork using a moving constraint that minimizes the center of mass distance between the group of Kap95 molecules and the nanopore occlusion. To confine the Kap95 proteins to the vicinity of the nanopore, a cylindrical occlusion (45 nm high, 90 nm diameter) of inert, 3 nm diameter beads was added on either side of the nanopore. This occlusion only interacts with Kap95. Following the pulling step, an equilibration simulation of 7.5×10^7^ steps (1.5*μ*s) was performed, where Kap95 molecules are constrained, such that the Nsp1 meshwork can homogenize before the 5×10^8^ steps (10 *μ*s) production run.

### Calculating the conductance of Nsp1-coated nanopores in the presence of Kap95

We calculated the conductance in a similar fashion to previous work^28,35,36^, relying on the Hall formula for the conductance of a nanopore, where the conductance originates from the serialized conductance of the pore region (|*z*| < 10 nm) and the access regions (10 < |*z*| < 40 nm):

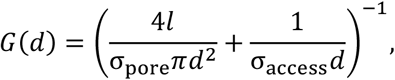

where *l* is the length of the nanopore channel (20 nm), *d* the pore diameter, and the conductance of the pore and access regions are given by *σ*_pore_ and *σ*_access_, respectively^53^. The conductance of the pore and access regions (where the latter is averaged over both sides of the pore) are calculated by integrating the local conductivity over the cylindrical volume that constitutes both regions:

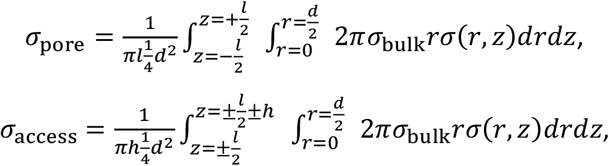

with *σ*_bulk_ the bulk conductivity of the 150 mM KCl solution (2.2 nS/nm). We incorporate the effect of local protein density on the conductance for both pore and access region by considering a spatially varying function that modulates the conductivity based on the local, time-averaged (using blocks of 10 ns length) protein density *ρ*:

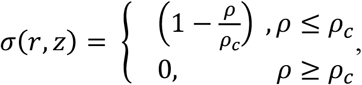

where we used a value of 85 mg/mL^35^.

### Residue contact analysis

We calculated the FG-pairing fraction using Python3.8 and the MDAnalysis package, version 1.9. For each trajectory frame, an upper triangular contact matrix *C* is calculated that describes all unique contacts between the F-residues in the FG-motifs. The value of element *C*_*ij*_ is set to 1 if the distance *d*_*ij*_ between the F-residues in FG-motifs *i* and *j* is smaller than a certain cut-off value, which is set at a value of 0.7 nm. The sum of the elements in the contact matrix *C* then yields the total number of F-F contacts for a given configuration (trajectory frame). A similar approach is taken for the Nsp1-Nsp1 contacts. We calculated the full contact matrices between all unique combinations of Nsp1-proteins inside the nanopore. This approach yields *N*_Nsp1_ (*N*_Nsp1_ − 1)/2 contact matrices (with *N*_Nsp1_ the number of Nsp1 proteins inside each nanopore), each containing the total number of contacts (defined as a distance *d*_*ij*_ < 1 nm) for each residue pair in the two Nsp1-molecules that occurred during the simulation. The average contact curve is then calculated by calculating a cumulative contact matrix for all Nsp1-Nsp1 pair, and normalizing this matrix against the length of the trajectory and the number of Nsp1 molecules. The matrix can be flattened to obtain the finanl Nsp1-Nsp1 contact curve. For Kap95-Nsp1 contacts, a similar approach is taken.

### Mapping of density and displacement data to axi-radial distributions

Axi-radial density distributions, averaged over time and azimuthal direction, were calculated using the `gmx_densmap` utility, where 3D density distributions are calculated by on a bin grid (0.5 nm bin size). Subsequently, an axi-radial coordinate system is defined, with the origin at the center-of-mass of the nanopore scaffold, which allows for a conversion of the 3D cartesian density distribution to an (azimuthally averaged) axi-radial density distribution. Axi-radial velocity maps were calculated in two steps: First, the center-of-mass displacement was calculated from the trajectories of individual Kap95 molecules, which by means of a central differences approach is converted into a velocity vector for each kap95 center of mass. The scalar velocity (speed) is then calculated from the vector norm of the velocity. Next, stochasticity from short-timescale movements was removed by applying a moving average with a window size of 250 frames to scalar velocities and coordinates. Next, the cartesian coordinates were converted to an axi-radial coordinate system, and a velocity map was calculated by spatially binning the velocities (0.5 nm bin size) and calculating the average value.

