## Supplementary information for "Transport receptor occupancy in Nuclear Pore Complex mimics"

[*]These authors contributed equally.

**1. 1/f noise comparison of bare pore vs Nsp1-coated pore**

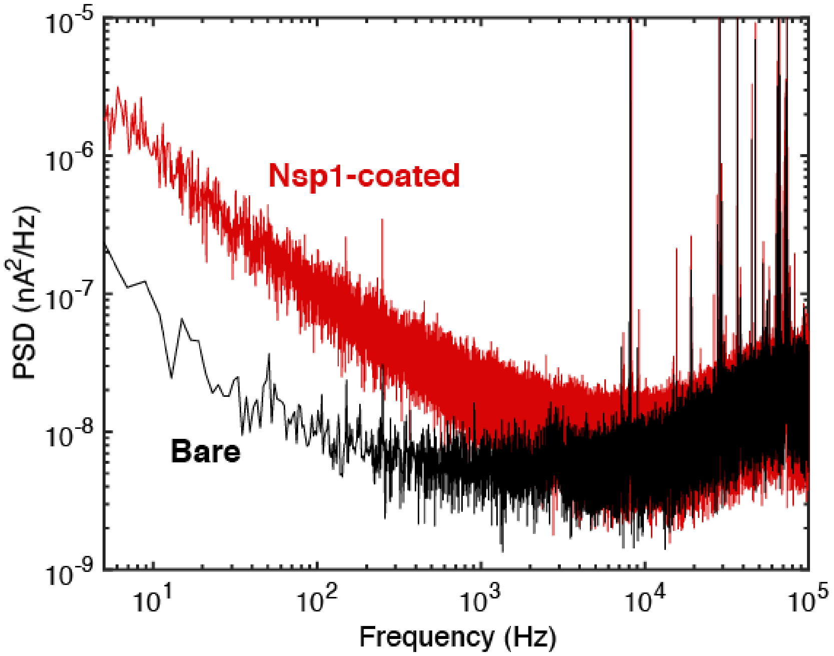

**Figure S1:** Current PSD spectra for a bare pore (black) vs a Nsp1-coated pore (red), illustrating the pro- nounced increase in 1/f noise upon coating the pore with Nsp1.

### 2. Additional traces of fast Kap95 translocations

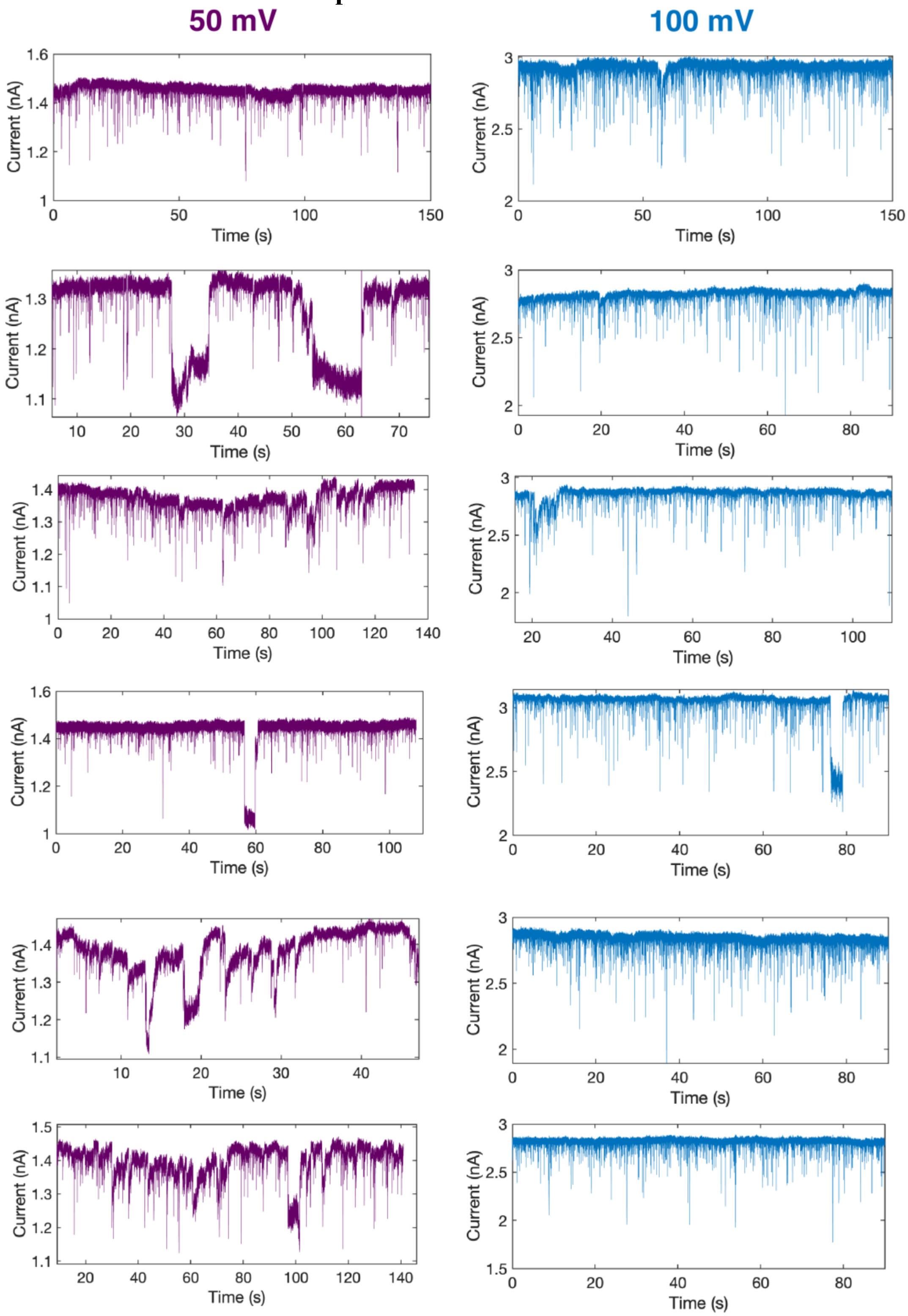

**Figure S2.** Additional current traces showing fast Kap95 translocation events through a Nsp1-coated pore under 50mV (left, purple) and 100mV (right, blue).

**3. SPR measurements of Nsp1-coated silica surfaces**

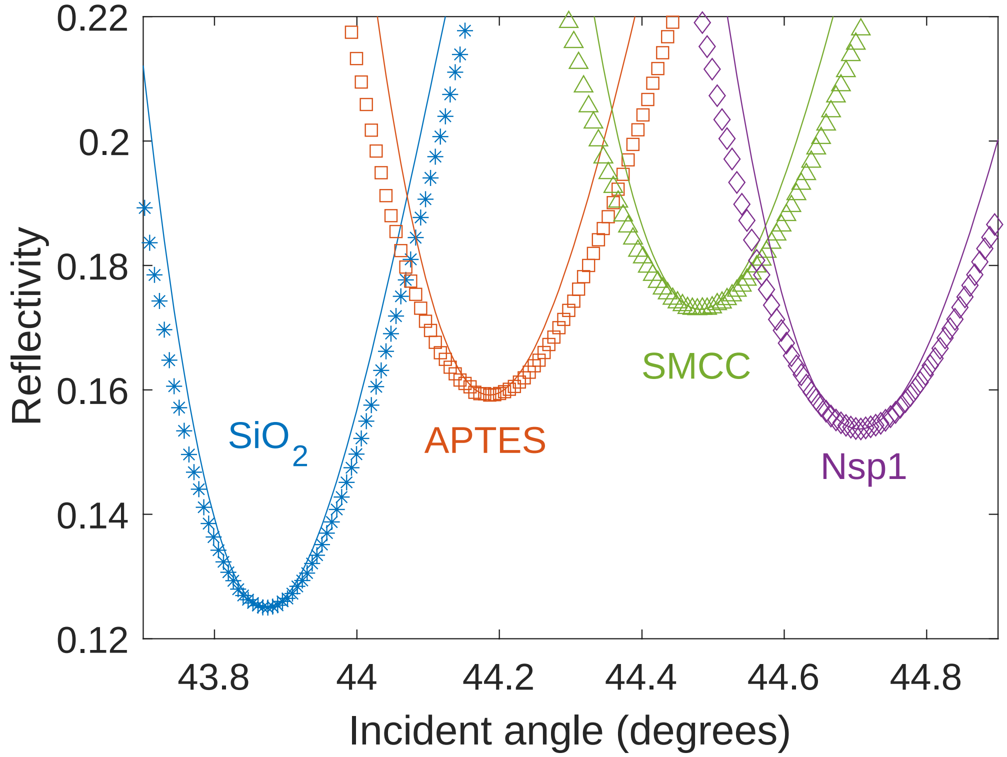

**Figure S3** SPR reflectivity measurement of SiO2 (blue), APTES (red), SMCC (green) and Nsp1 (magenta) layers.

### 4. Residue-scale 1-BPA forcefield with inclusion of cation-pi interactions

The one-bead-per amino acid (1-BPA) model distinguishes between all twenty amino acids based on their hydrophobicity, charge and backbone stiffness[1–4]. In the current work, a modified version termed 1-BPA-CP is used, where the hydrophobicity scale values of charged residues are modified[4], and where the Lennard-Jones-type interaction between aromatic and cationic residues is strengthened[5].

Non-bonded interactions between amino acid residues comprise hydrophobic interactions, charged interactions and cation-pi interactions. Hydrophobic interactions ($\Phi_{\mathrm{HP}}$) are described using a shifted 8-6 Lennard Jones-type potential:

$$\Phi_{\mathrm{HP}} = \begin{cases} \epsilon_{\mathrm{rep}}\left(\frac{\sigma}{r}\right)^8 - \epsilon_{ij}\left[\frac{4}{3}\left(\frac{\sigma}{r}\right)^6 - \frac{1}{3}\right], & r \leq \sigma \\ \left(\epsilon_{\mathrm{rep}} - \epsilon_{ij}\right)\left(\frac{\sigma}{r}\right)^8, & \sigma \leq r \end{cases} \quad (1)$$

where the interaction strengths follow from a repulsive component $\epsilon_{\mathrm{rep}}$ = 10 kJ/mol, and an attractive component $\epsilon_{ij} = 13 \cdot \sqrt{(\epsilon_i \epsilon_j)^{\alpha}}$ kJ/mol that depends on the individual hydrophobicity-value $\epsilon_i$ (Table S1) and a global fitting parameter $\alpha$=0.27. Electrostatic interactions ($\Phi_{\mathrm{EL}}$) follow from a modified Coloumb law:

$$\Phi_{\mathrm{EL}} = \frac{q_i q_j}{4\pi\epsilon_0 \epsilon_{\mathrm{r}}(r) r} \exp\left(-\kappa r\right), (2)$$

where a salt concentration of 150 mM KCl is implicitly accounted for via the Debye screening term (with $\kappa = 1.27$ nm$^{-1}$) and a distance-dependent dielectric constant $\epsilon_r$. This constant accounts for local solvent re-structuring and is given as follows:

$$\epsilon_r(r) = 80\left[1 - \frac{r^2}{z^2}\frac{e^{\frac{r}{z}}}{\left(e^{\frac{r}{z}}-1\right)^2}\right], (3)$$

where $z = 0.25$ nm.

Rather than using the 1-BPA hydrophobic potential, residue pairs that engage in cation-pi interactions (e.g., F, Y or W with R or K) interact through an 8-6 Lennard-Jones potential $\phi_{\mathrm{cp,ij}}$:

$$\phi_{\mathrm{cp,ij}}(r) = \varepsilon_{\mathrm{cp,ij}}\left[3\left(\frac{r_{\mathrm{m}}}{r}\right)^8 - 4\left(\frac{r_{\mathrm{m}}}{r}\right)^6\right], (4)$$

where $\varepsilon_{\mathrm{cp,ij}}$ is the pair-dependent interaction energy and $r_{\mathrm{m}}$ the equilibrium distance, set at 0.45 nm. The interaction parameters between all residue pairs are provided in Table S2.

**5. Parametrization of binding site strength in the single-residue yeast Kap95 model**

We assigned a specific interaction strength $\epsilon_{\mathrm{BS,FG}}$ (rather than $\epsilon_{\mathrm{ij}}$) between binding site regions on Kap95 and any FG-motif residue (Figure S5a, Tables S3-4). An experimentally-determined lower limit for the dissociation constant between Kap95 and a highly-charged Nsp1-segment ('FSFG-K', Fig. S5b) of 36.1 μM exists: by calculating the dissociation constant between Kap95 and FSFG-K from our simulations for varying $\epsilon_{\mathrm{BS,FG}}$ (Figure S5c), we find that this lower bound can be reproduced for $\epsilon_{\mathrm{BS,FG}}$=13.88 kJ/mol, which is correspondingly an upper bound to the value of $\epsilon_{\mathrm{BS,FG}}$. We observe that the association between Kap95 and FSFG-K is characterized by frequent, multivalent and transient binding events between binding sites and FG-motifs (Figure S5d) for a range of $\epsilon_{\mathrm{BS,FG}}$-values. This is in line with the binding mechanism as described in earlier work[6,7].

To assess the transport properties of our Kap95 protein independently from the Nsp1 nanopore system, we employed a nanopore coated with an artificial FG-Nup 'NupX' that we studied in earlier work[8]. Kap95 is known to translocate through NupX-coated nanopores. We used a qualitative criterion that postulates that Kap95 should localize inside the pore meshwork (in absence of any external bias voltage or enzymatic activity with RanGTP) and maintain the largest possible mobility to refine our value for $\epsilon_{\mathrm{BS,FG}}$. As shown in Figure S6, we find that for a range of $\epsilon_{\mathrm{BS,FG}}$-values, Kap95 localizes inside the NupX-pore, with a lower bound at 13.7 kJ/mol. When considering the mobility of the Kap95 molecule inside the nanopore meshwork (Figure S6), highly valent and transient binding (Figure S5d) and the experimental $K_{\mathrm{D}}$-value, we find that a value of $\epsilon_{\mathrm{BS,FG}}$13.75 kJ/mol best describes the desired behavior.

### 6. Structure and genetic alignment of yeast and mouse importinβ

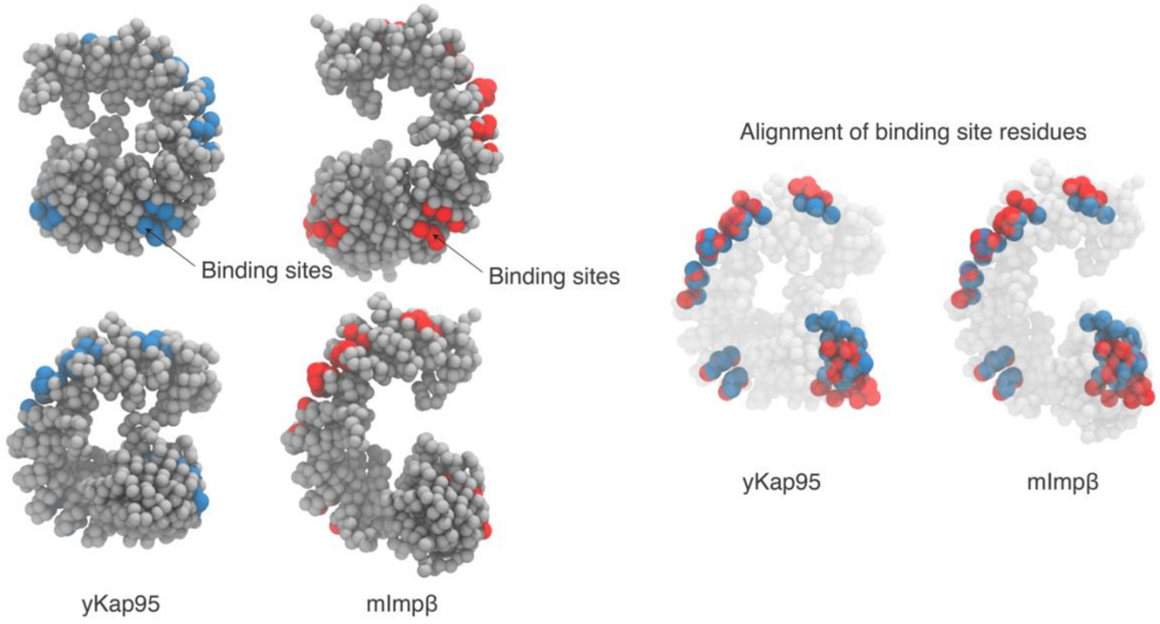

**Figure S4:** Mapping the binding site residues identified by Isgro and Schulten[9] in mImpβ to yKap95 by means of a sequence alignment leads to good structural conservation of the binding site positions. Left: structural alignment highlights the similarity of yKap95 and mImpβ and their FG-motif-binding site regions. Right: structural alignment of binding site residues, projected onto either yKap95 or mImpβ. Binding site residues in mImpβ are labeled in red, evolutionary conserved counterparts in yKap95 in blue, demonstrating a good spatial overlap between both structural models.

### 7. Parametrization of interactions between FG-motifs and Kap95 binding site regions

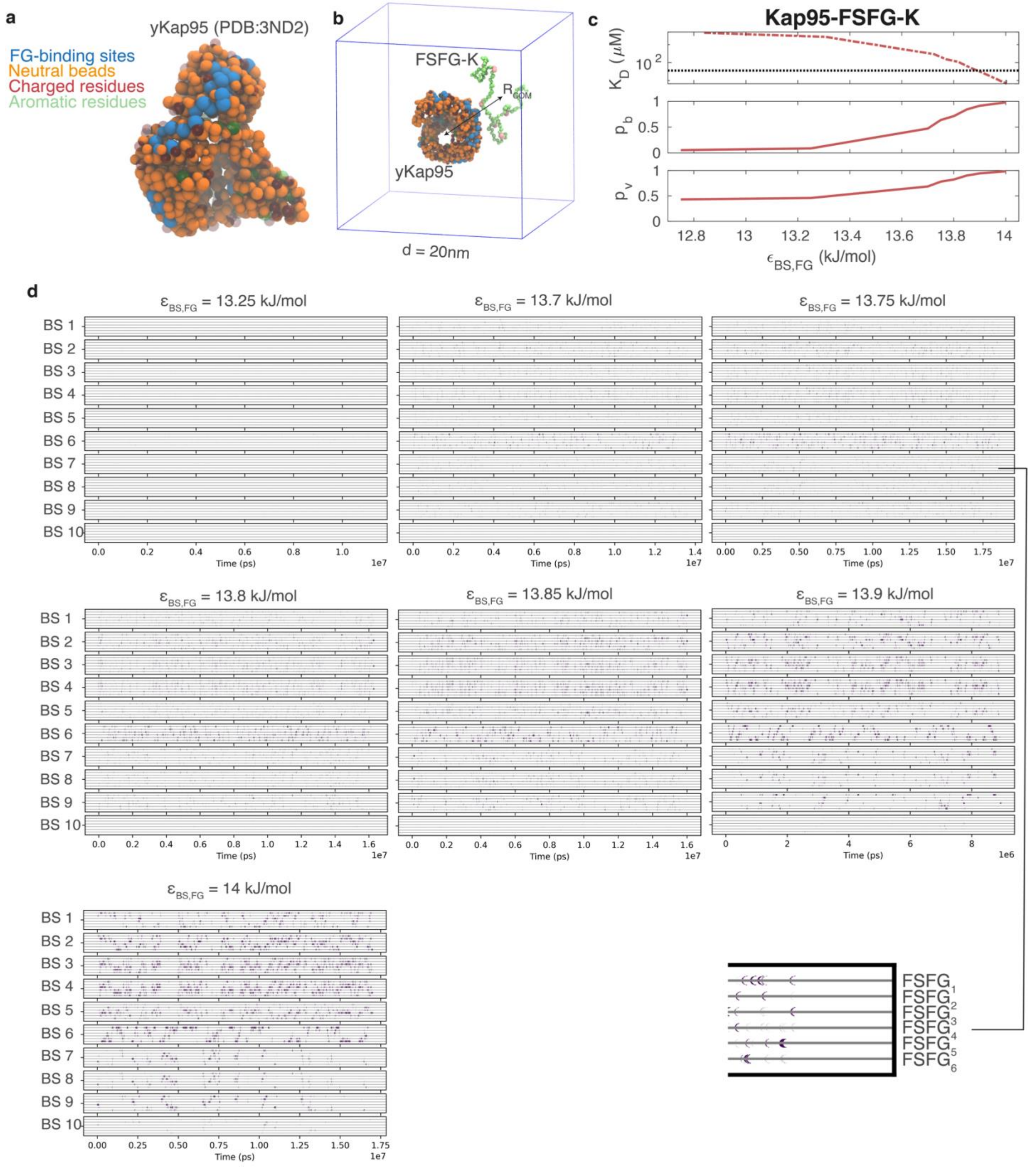

**Figure S5:** Parametrizing the coarse-grained model for yeast Kap95**. a:** Schematic depiction of the residue-scale Kap95 model, with key bead types highlighted. **b:** We parametrized the strength of the binding site-FG-motif interactions (blue on Kap95, pink on FSFG-K, respectively) by reproducing the dissociation constant $K_{\mathrm{D}}$ between Kap95 and a highly-charged Nsp1-segment 'FSFG-K' using the sub-volume method. $P_{\mathrm{b}}$ denotes the fraction of bound states and $P_{\mathrm{v}}$ indicates interacting states where the center-of-mass distance of the proteins falls within the radius of the interaction sub-volume. **c:** Values of $K_{\mathrm{D}}$ (top), $P_{\mathrm{b}}$ (middle) and $P_{\mathrm{v}}$ (bottom) as a function of FG-motif-binding site strength $\epsilon_{\mathrm{BS,FG}}$. An intersect with the experimentally-determined lower bound for the dissociation constant of FSFG-K and Kap95 (36.1 μM) is obtained at $\epsilon_{\mathrm{BS,FG}}$=13.88 kJ/mol. **d:** Analysis of contacts in time between individual Kap95 binding sites and FG-motifs in the FSFG-K protein, for varying binding site strengths. For each of the 10 binding sites on Kap95 (vertically stacked graphs), timestamps where a distance of < 0.7 nm between any binding site residue and an FG-motif occurs are indicated in purple. Within each binding site time-series graph, the contacts of Kap95 in time with of individual FG-motifs is plotted on grey horizontal axes (see inset for close-up). For binding sites 1, 5, 6 and 10, additional residues were incorporated into the binding site based on their geographic proximity to evolutionarily conserved binding site residues[9] such that the different binding sites exhibit more similar binding statistics. Binding site 10 shows a significantly lower number of contacts due to its positioning on the inner surface of Kap95.

### 8. Refining the binding site strength based on simulations of Kap95 inside NupX-coated nanopores

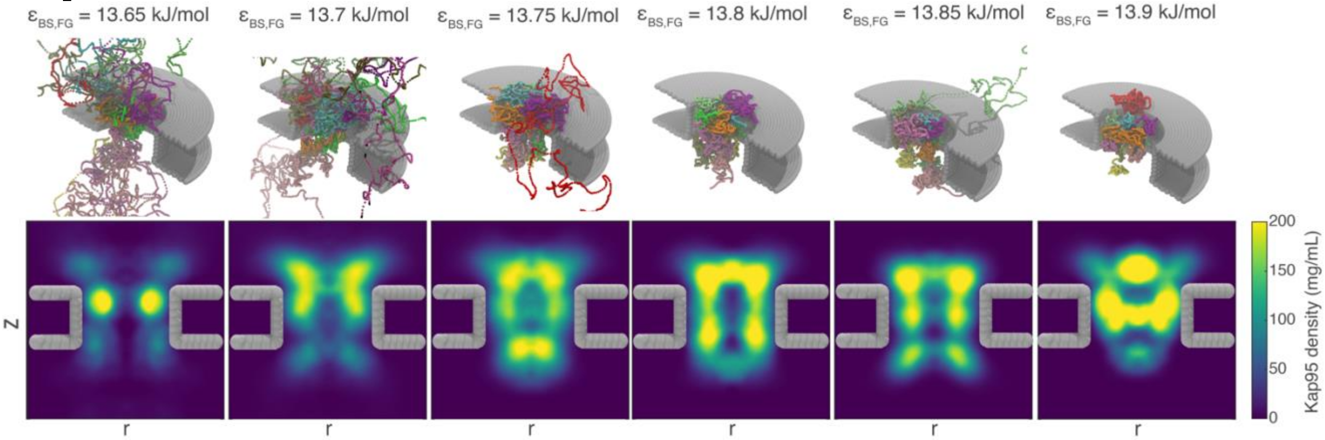

**Figure S6** Using a coarse-grained simulation of 10 copies of Kap95 inside a 30 nm diameter pore coated with the artificial FG-Nup 'NupX', we identified solutions for $\epsilon_{\mathrm{BS,FG}}$ that demonstrate adsorption of Kap95 into the pore lumen, while allowing the largest degree of mobility for Kap95. **Top row**: Center-of-mass trajectories of individual Kap95 molecules (smoothened and plotted in different colors for each Kap95 molecule) for varying binding site strength. **Bottom row:** Axi-radial time-averaged density distributions of Kap95. Increasing the binding site strength beyond 13.75 kJ/mol leads to a reduced mobility of Kap95 within the pore lumen. Based on the combined results in terms of $K_{\mathrm{D}}$ (Fig. S5c), highly valent and transient binding (Fig. S5d), and maximum mobility of Kap95 while being retained in the NupX-meshwork, we selected $\epsilon_{\mathrm{BS,FG}}$=13.75 kJ/mol as the optimal value for the strength of the FG-motif-binding site interaction.

### 9. Convergence analysis of binding affinity calculations

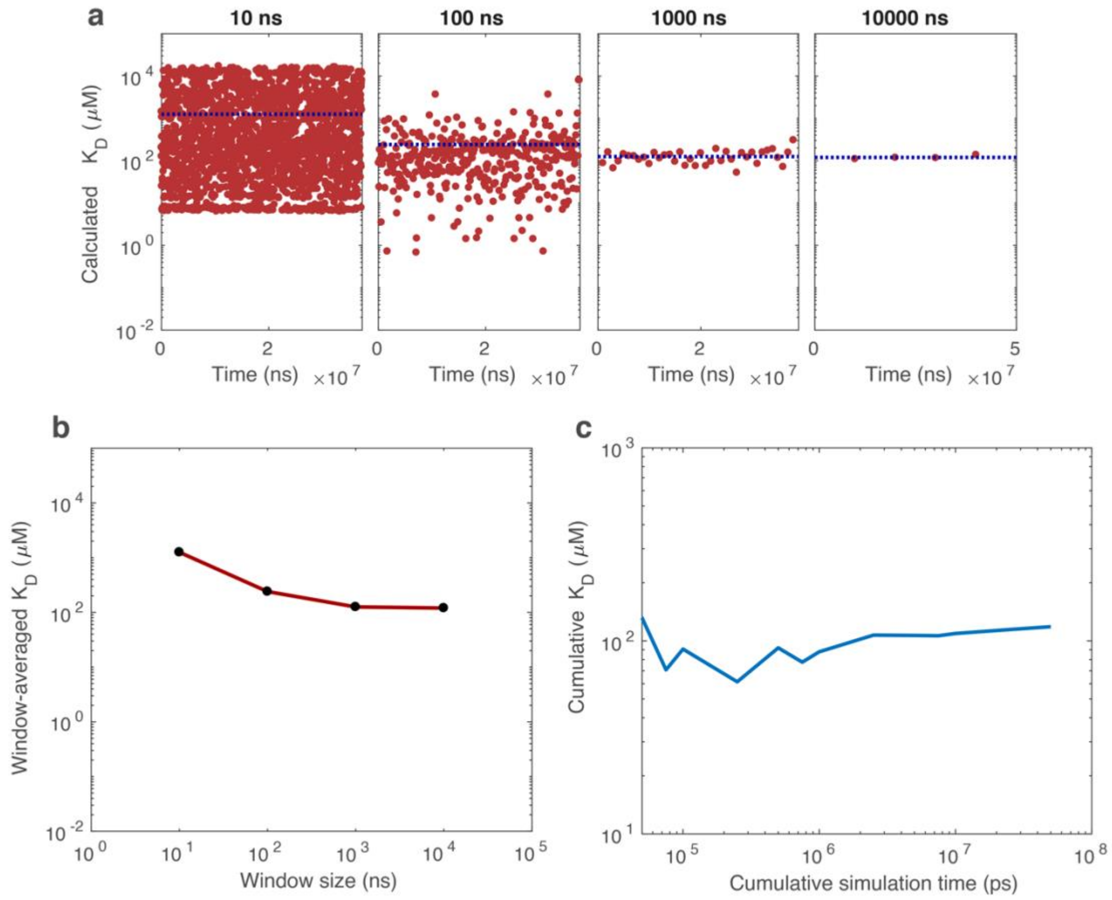

**Figure S7:** Convergence analysis of $K_D$-calculations. **a:** $K_{\mathrm{D}}$-values calculated from trajectory windows with varying length. Each mark indicates the $K_{\mathrm{D}}$-value calculated from a single window; the blue dotted line indicates the average $K_{\mathrm{D}}$-value calculated from all marks. The variation in $K_{\mathrm{D}}$-values around the mean becomes reasonably small at a window size of 1 microsecond (1000 ns). **b:** Average $K_{\mathrm{D}}$-value as a function of window size, corresponding to the blue lines in panel a. Increasing the window size beyond one microsecond (1000 ns) does not lead to a further change in the average $K_{\mathrm{D}}$-value. **c:** Estimate of $K_{\mathrm{D}}$ as a function of the total simulation time. Consistent with panels a, b and results from Lopez *et al.*[10], we find that 1-2 microseconds (approx. $10^4$ configurations) are necessary to obtain statistical convergence of the $K_{\mathrm{D}}$ -value.

**10. Average binding statistics between Kap95 and Nsp1**

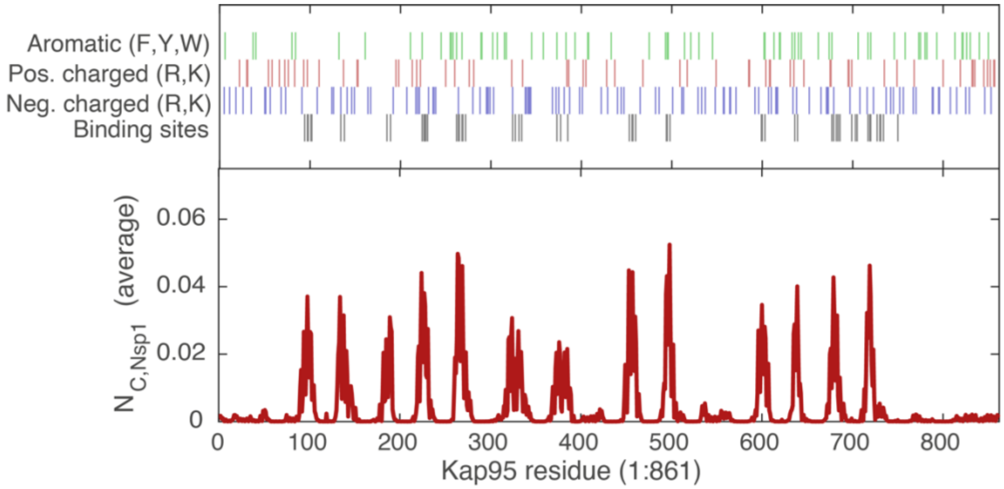

**Figure S8:** Time-averaged number of contacts between Kap95 and Nsp1-proteins inside Nsp1-coated nanopores for a separation <1.0 nm. The interactions between Kap95 and Nsp1 are driven by the binding site regions on Kap95 and FG-motifs in Nsp1. Since only exposed residues constitute a binding site on the surface of Kap95, the number of peaks in the lower panel does not necessarily correspond to the number of binding sites.

**11. Persistence length of Nsp1 as a function of Kap95 occupancy**

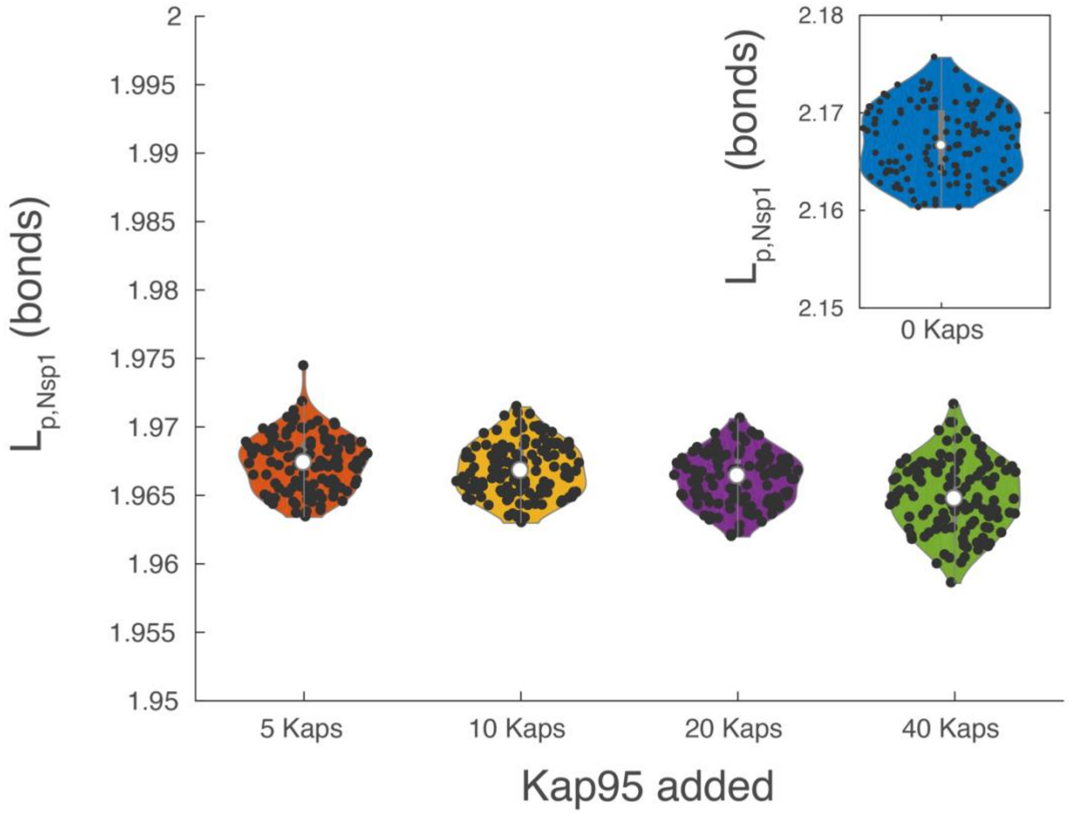

**Figure S9** Average persistence length (in number of bonds) of the Nsp1-proteins inside the Nsp1-coated nanopores, as a function of the number of added Kap95 molecules. A slight (~10%) reduction in persistence length of the individual Nsp1 molecules is observed as Kap95 is added, where the size of the effect does not depend on Kap95 concentration.

### 12. Normalized hydrophobicity values used within the 1-BPA-CP model

| **AA** | A | R* | N | D* | C | Q | E* | G | H | I | L | K* | M | F | P | S | T | W | Y | V |
|---|---|---|---|---|---|---|---|---|---|---|---|---|---|---|---|---|---|---|---|---|
| $\varepsilon_i$ | 0.7 | 0.005 | 0.33 | 0.005 | 0.68 | 0.64 | 0.005 | 0.41 | 0.53 | 0.98 | 1 | 0.005 | 0.78 | 1 | 0.65 | 0.45 | 0.51 | 0.96 | 0.82 | 0.94 |

**Table S1:** Interaction parameters used in the 1-BPA-CP model used throughout this work. *The hydrophobicity values of charged residues have been slightly increased in line with recent work[4].

**13. Cation-pi interaction strengths within the 1-BPA-CP forcefield**

| Aromatic residue | Cationic residue | $\varepsilon_{cp,ij}$ (kJ/mol) |
|---|---|---|
| F | R | 4.3 |
| F | K | 1.79 |
| Y | R | 5.0 |
| Y | K | 3.13 |
| W | R | 6.7 |
| W | K | 4.26 |

**Table S2:** Interaction parameters $\varepsilon_{cp,ij}$ between aromatic and cationic amino acids. Values adapted from Jafarinia *et al*[5].

**14. Particle types used in coarse-grained modeling**

| **Particle type** | **Consists of:** | **$\sigma$ (nm)** |
|---|---|---|
| **AA** | All residues not involved in cation-pi (RK/FYW) or binding site (B-RK, B-FYW, F1, G1) interactions | **0.6** |
| **RK** | R or K-residues that are not part of a Kap95 binding site | **0.6** |
| **FYW** | F, Y or W residues that are not part of a Kap95 binding site or FG-motif (in case of F). | **0.6 (hydrophobic) or 0.45 (cation-pi)** |
| **B-RK** | R or K-residues that are part of a Kap95 binding site | **0.6 or 0.45** |
| **B-FYW** | F, Y or W residues that are part of a Kap95 binding site | **0.6 or 0.45** |
| **F1** | F-residue within an FG-motif | **0.6** |
| **G1** | G-residue within an FG-motif | **0.6** |
| **WALL** | Sterically beads that constitute the nanopore scaffold | **3** |
| **BOUNDS** | Sterically inert beads that confine only Kap95 to a cylindrical region on either side of the nanopore membrane | **3** |

**Table S3:** Overview of interaction types

**15. Interaction table for different particle types used in coarse-grained modeling**

| | AA | RK | FYW | B-RK | B-FYW | F1 | G1 | WALL | BOUNDS |
|---|---|---|---|---|---|---|---|---|---|
| AA | 1BPA | | | | | | | | |
| RK | 1BPA | 1BPA | | | | | | | |
| FYW | 1BPA | CP | 1BPA | | | | | | |
| B-RK | 1BPA | 1BPA | CP | 1BPA | | | | | |
| B-FYW | 1BPA | CP | 1BPA | CP | 1BPA | | | | |
| F1 | 1BPA | CP | 1BPA | BS | BS | 1BPA | | | |
| G1 | 1BPA | 1BPA | 1BPA | BS | BS | 1BPA | 1BPA | | |
| WALL | 1BPA* | 1BPA* | 1BPA* | 1BPA* | 1BPA* | 1BPA* | 1BPA* | EXCL | |
| BOUNDS | EXCL | EXCL | EXCL | 1BPA* | 1BPA* | EXCL | EXCL | EXCL | EXCL |

**Table S4:** Interaction parameters between coarse-grained beads. 1BPA denotes the hydrophobic potentials as indicated in equation S1, CP denotes residues interacting via cation-pi interactions (equation S4), BS denotes the hydrophobic potential (equation S1) with the parametrized binding site strength, 1BPA* indicates the hydrophobic potential (equation S1) with only a repulsive term and with one particle having a diameter of 3 nm, and EXCL indicates excluded interactions that are not calculated.

**16. Binding site residues in yeast Kap95 obtained from an evolutionary and structural alignment analysis**

| Binding site # | Residue numbers |
|---|---|
| 1 | **<u>94</u>**, 97, **<u>98</u>**, 101, 102, 134, 138 |
| 2 | 185, 189, 224, 227, 228 |
| 3 | 226, 230, 262, 269 |
| 4 | 264, 265, 268, 269, 272, 324 |
| 5 | 327, 331, 373, 385, **<u>334</u>**, **<u>377</u>**, 385 |
| 6 | 453, 456, 457, 460, 494, 495, **<u>498</u>** |
| 7 | 599, 600, 603, 636, 639 |
| 8 | 603, 677, 684 |
| 9 | 679, 682, 686, 717, 719, 720 |
| 10* | 699, 703, 705, 727, **<u>730</u>**, 731, **<u>734</u>**, 750 |

**Table S5:** Binding site residues in a single-residue coarse-grained Kap95 model. We identified the binding site residues in yeast Kap95 based on a sequence alignment to mouse impβ, for which the binding site residues were determined in earlier work[9] . Additional residues (bold, underlined) were added to certain sites to enhance the accessibility of the binding site to FG-motifs. *Binding site 10 is oriented inwards to the convex surface of Kap95 and was found to not participate in binding with FG-motifs as well as other sites[9].

**17. Overview of all simulations in this work**

| System | System size (no. of beads) | Number of simulations | Forcefield | Timestep (ps) | Simulation time (steps) | Temperature (K) |
|---|---|---|---|---|---|---|
| FSFG-K with Kap95 | 964 | 20 per value of $\epsilon_{\mathrm{BS,FG}}$ | 1-BPA-CP | 0.02 | 2000000000 (cumulative) | 300 |
| NupX-nanopore (30nm) with 10 x Kap95 | 32444 | 6 | 1-BPA-CP | 0.02 | 100000000 | 300 |
| Nsp1 nanopore (55nm), equilibration | 82811 | 1 | 1-BPA-CP | 0.02 | 25000000 | 300 |
| Nsp1 nanopore (55nm) | 82811 | 1 | 1-BPA-CP | 0.02 | 500000000 | 300 |
| Nsp1 nanopore (55nm) with 5x Kap95, equilibration | 112026 | 1 | 1-BPA-CP | 0.02 | 75000000 | 300 |
| Nsp1 nanopore (55nm) with 5x Kap95 | 112026 | 1 | 1-BPA-CP | 0.02 | 500000000 | 300 |
| Nsp1 nanopore (55nm) with 10x Kap95, equilibration | 116331 | 1 | 1-BPA-CP | 0.02 | 75000000 | 300 |
| Nsp1 nanopore (55nm) with 10x Kap95 | 116331 | 1 | 1-BPA-CP | 0.02 | 500000000 | 300 |
| Nsp1 nanopore (55nm) with | 124941 | 1 | 1-BPA-CP | 0.02 | 75000000 | 300 |

| | | | | | | |
|---|---|---|---|---|---|---|
| 20x Kap95, equilibration | | | | | | |
| Nsp1 nanopore (55nm) with 20x Kap95 | 124941 | 1 | 1-BPA-CP | 0.02 | 500000000 | 300 |
| Nsp1 nanopore (55nm) with 40x Kap95, equilibration | 142161 | 1 | 1-BPA-CP | 0.02 | 75000000 | 300 |
| Nsp1 nanopore (55nm) with 40x Kap95 | 142161 | 1 | 1-BPA-CP | 0.02 | 500000000 | 300 |

**Table S6** Overview of all simulations in this work.
